# Self-incompatibility phenotypes of SRK mutants can be predicted with high accuracy

**DOI:** 10.1101/2024.04.10.588956

**Authors:** Masaya Yamamoto, Shotaro Ohtake, Akihisa Shinosawa, Matsuyuki Shirota, Yuki Mitsui, Hiroyasu Kitashiba

**Affiliations:** Graduate School of Agricultural Science, Tohoku University, 468-1 Aramaki Aza Aoba, Aoba-ku, Sendai, Miyagi 980-8572, Japan; NODAI Genome Research Center, Tokyo University of Agriculture, 1-1-1 Sakuragaoka, Setagaya-ku, Tokyo, 156-8502, Japan; Graduate School of Medicine, Tohoku University, 2-1 Seiryo-machi, Aoba-ku, Sendai, Miyagi 980-8575, Japan; Graduate School of Agricultural Science, Tokyo University of Agriculture, 1237 Funako, Atsugi, Kanagawa 243-0034, Japan

## Abstract

Only very limited information is available on why some non-synonymous variants severely alter gene function while others have no effect. To identify the characteristic features of mutations that strongly influence gene function, this study focused on *S-locus receptor kinase*, *SRK*, which encodes a highly polymorphic receptor kinase expressed in stigma papillary cells that underlies a female determinant of self-incompatibility in Brassicaceae. A set of 299 *Arabidopsis thaliana* transformants expressing mutated *SRKb* from *A. lyrata* was constructed and analyzed to determine the genotype and self-incompatibility phenotype of each transformant. Almost all the transformants showing the self-incompatibility defect contained mutations in AlSRKb that altered localization to the plasma membrane. The observed mutations occurred in amino acid residues that were highly conserved across *S* haplotypes and whose predicted locations were in the interior of the protein. These mutations were likely to underlie the self-incompatibility defect as they caused significant changes to amino acid properties. Such findings suggested that mutations causing the self-incompatibility defect were more likely to result from changes to AlSRKb biosynthesis than from loss of function. In addition, this study showed the RandomForest and Extreme Gradient Boosting methods could predict self-incompatibility phenotypes of SRK mutants with high accuracy.

## INTRODUCTION

Errors in DNA replication that result in a change in the DNA sequence produce nucleotide variation in living organisms. Although non-synonymous variants may cause genes to lose their function or develop new ones, some have no effect on the functions of genes. Advances in sequencing technology over the past two decades have revealed the full picture of DNA polymorphisms across a genome, enabling the construction of genome-wide association study (GWAS) platforms in many organisms. Identifying the causative single nucleotide variants (SNVs) within the set of genome-wide DNA polymorphisms is a crucial step in genetic analysis, but also the most difficult. Therefore, identifying the characteristics of non-synonymous variants that affect gene function will be useful for finding causative SNVs in GWAS and quantitative trait locus (QTL) mapping analyses, and will also contribute to resolving the pressing challenges facing agriculture and human healthcare.

Genes with high allelic diversity are suitable for determining the characteristics of non-synonymous variants with large influences on gene function, especially when each allele has a different effect. We therefore focused on the self-incompatibility (SI) mechanism in Angiosperms. SI enables plants to avoid self-fertilization; by facilitating cross-fertilization, it avoids inbreeding depression and maintains genetic variation. About 40% of Angiosperm families show SI (Barrett, 2002; Igić and Kohn, 2006; McCubbin and Kao, 2000). SI is usually determined by a single locus, the *S* locus, which contains several genes, and forms a distinctive haplotype, called the *S* haplotype (Silva and Goring 2001). Population genetic theory predicts that many *S* haplotypes should be maintained, given that individuals possessing rare *S* haplotypes have more mating opportunities than those carrying common haplotypes (Wright, 1939, Schierup 1998). Consistent with this, more than 50 *S* haplotypes, each of which shows different self-recognition activity, are known from cultivated *Brassica* species (Oikawa et al. 2011, Yamamoto et al. 2023).

SI in the Brassicaceae is genetically controlled by two tightly linked, highly polymorphic genes within the *S* locus. *S-locus receptor kinase* (*SRK*) encodes a plasma membrane-localized receptor kinase expressed in stigmatic papillae cells (Stein et al. 1991, Takasaki et al. 2000) and *S-locus cysteine-rich protein*/*S-locus protein 11* (*SCR*/*SP11*, hereafter referred to as *SCR*) encodes a cysteine-rich peptide ligand of SRK displayed at the pollen surface (Schopfer et al. 1999, Takayama et al. 2000). SRK can only interact with SCR of the same *S* haplotype. This allele-specific SRK-SCR interaction activates the SI response, inhibiting pollen germination and growth of the pollen tube on the stigma surface. Functional diversification of *S* haplotypes is achieved by multiple amino acid polymorphisms in SRK and SCR. The mature region of SCR shows extraordinary diversity, including frequent insertions and deletions (Sato et al. 2002). Although similarity of full-length SRK is relatively high compared with SCR (Sato et al. 2002), three hypervariable regions (hv I, II, and III) are found in the receptor domains of SRK in the *S* haplotypes in *Brassica* and *Arabidopsis* species (Kusaba et al. 1997, Nishio and Kusaba, 2000, Castric and Vekemans, 2007). Structural analysis of the SRK-SCR complex suggests that amino acid residues in the hv regions of SRK are involved in the allele-specific SRK-SCR interaction (Ma et al. 2016, Murase et al. 2020). Since SRK shows high diversity in allelic sequences as well as a specific recognition activity, we considered it a suitable protein model for unraveling the characteristics of non-synonymous variants with influences on gene function.

The model plant *Arabidopsis thaliana* is a useful experimental tool for analyzing the effects of mutations on gene function because it can be transformed simply and efficiently. Although *A. thaliana* shows self-compatibility (SC), as its *S* locus contains either a nonfunctional *SCR* gene or nonfunctional *SRK* and *SCR* genes (Kusaba et al., 2001; Shimizu et al., 2008; Tang et al., 2007; Tsuchimatsu et al., 2010, 2017), it can be rendered SI by introducing the *SRK* and *SCR* genes of *Sb*, also known as the *S20* haplotype, from *Arabidopsis lyrata*, a closely-related SI species (Nasrallah et al. 2002, 2004). We examined the effects of mutations in *A. lyrata* SRKb (AlSRKb) on SI activity in *A. thaliana* transformants expressing AlSRKb containing randomly introduced mutations. We determined the SI phenotypes and genotypes of 299 *A. thaliana* transformants, and also measured the levels of AlSRKb expression and localization in the plasma membrane in 139 transformants. Almost all mutations in AlSRKb that caused SI defects also affected localization to the plasma membrane. Mutations in amino acid residues that were highly conserved across *S* haplotypes and located within the interior of the AlSRKb molecule resulted in significant changes to amino acid properties, which were associated with SI defects. In addition, we found we could predict the SI phenotypes of *A. thaliana* transformants with high accuracy (about 80%) by using the RandomForest and Extreme Gradient Boosting methods.

## RESULTS

### SI phenotypes of *A. thaliana* transformants expressing mutated AlSRKb-FLAG

To elucidate the effects of mutations in AlSRKb on SI of transgenic *A. thaliana*, we used the error-prone PCR method to mutate the coding region of the receptor domain of *AlSRKb.* We constructed a set of *A. thaliana* transformants expressing mutated C-terminal FLAG-tagged AlSRKb and AlSCRb (hereafter, mutated AlSRKb) (Figure 1A). This process generated 342 transformants. The SI phenotypes of 299 transformants were examined by manual pollination of flower buds immediately prior to opening (position-1) with pollen expressing AlSCRb. “Strong” and “weak” SI responses were defined by the growth of < 10 and 10 to 29 pollen tubes per stigma, respectively; the growth of > 29 pollen tubes per stigma was considered as an SC response. Of the 299 transformants tested, 105 transformants showed a strong SI response [SI], three showed a weak SI response [wSI], and 191 showed an SC response [SC] (Figure 1B; Table 1).

**Figure 1.**
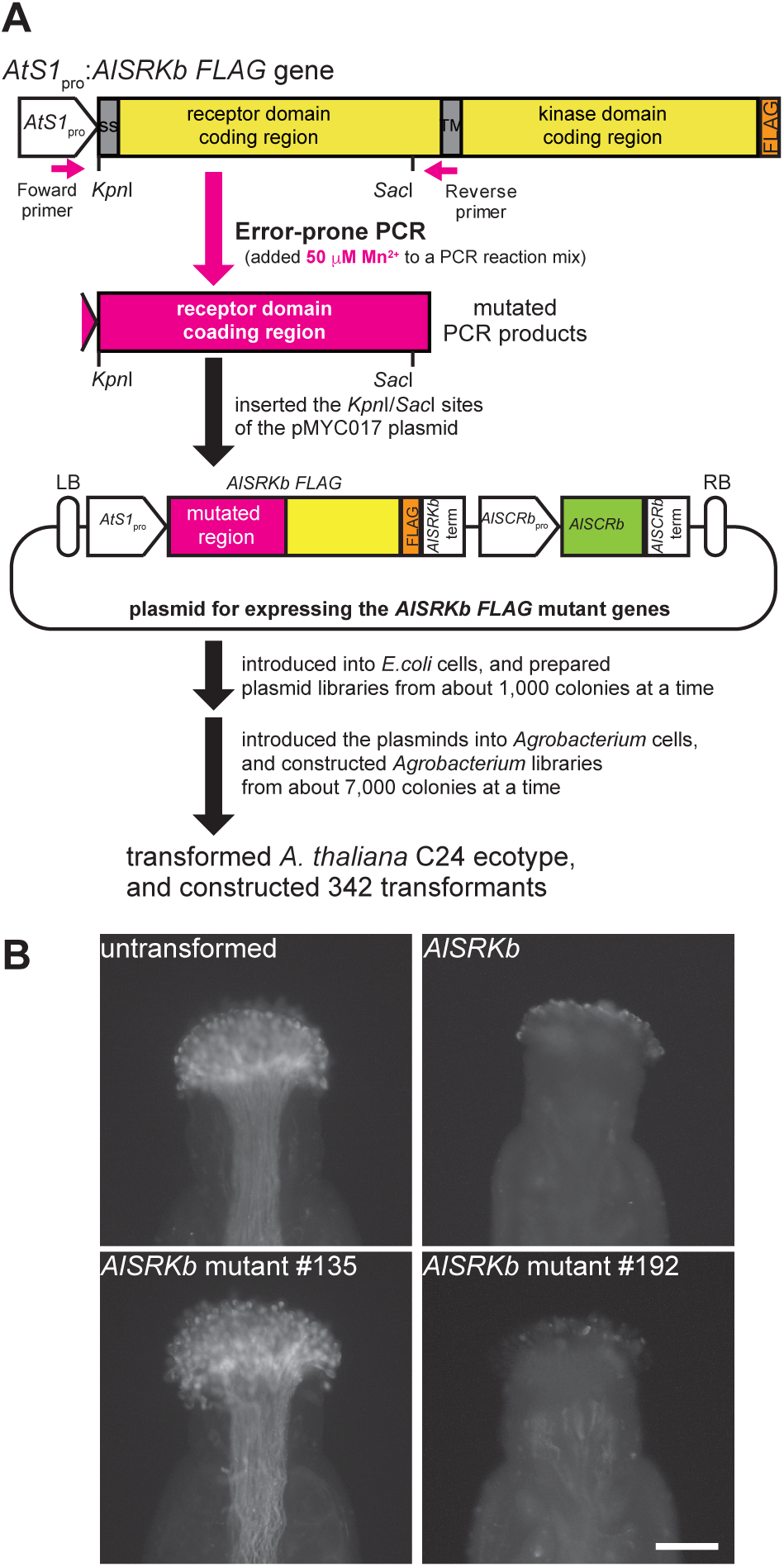
**Construction and phenotypic analysis of *A. thaliana* transformants expressing mutated *AlSRKb-FLAG*.** **(A)** Schematic diagram showing construction of *A. thaliana* transformants expressing *AlSRKb-FLAG* genes containing random mutations introduced using error-prone PCR. **(B)** Microscopic observations of pollinated stigmas of *A. thaliana* showing AlSCRb-expressing pollen stained with aniline blue. Untransformed: pollinated stigma from an untransformed *A. thaliana* plant; *AlSRKb*: pollinated stigma from an *A. thaliana* transformant expressing wild-type *AlSRKb-FLAG* + *AlSCRb*; *AlSRKb* mutant #135: pollinated stigma from *A. thaliana* transformant #135 expressing mutated *AlSRKb-FLAG* + *AlSCRb*; *AlSRKb* mutant #192: pollinated stigma from *A. thaliana* transformant #192 expressing mutated *AlSRKb-FLAG* + *AlSCRb*. The full results of all the analyzed phenotypes are listed in Supplemental Data Set 1. Scale bar = 100 μm.

**Table 1.** Summary of SI phenotypes of *Arabidopsis thaliana* transformants expressing mutated forms of the *Arabidopsis lyrata* gene *SRKb*.

| Mutation type | No. of self-pollen tubes per stigma<sup>a</sup> |  |  | Total |
| --- | --- | --- | --- | --- |
|  | <10 [SI] | 10–29 [wSI] | >29 [SC] |  |
| Non-synonymous mutation(s)<br>(and synonymous mutation(s)) | 41 | 1 | 53 | 95 |
| No start codon, frameshift, or<br>nonsense mutation(s) | 0 | 0 | 90 | 90 |
| Lack of *AlSRKb* coding *S* domain | 0 | 0 | 31 | 31 |
| Synonymous mutation(s) only | 16 | 0 | 2 | 18 |
| No mutation | 35 | 0 | 0 | 35 |
| Multi-insertion | 13 | 2 | 15 | 30 |
| Total | 105 | 3 | 191 | 299 |

<sup>a</sup>The number of pollen tubes per stigma was counted 2 h after self-pollination. Values of < 10, 10–29, and > 29 indicate strong SI [SI], weak SI [wSI], and compatible responses [SC], respectively.

The DNA sequences of the mutated region of *AlSRKb* were determined by the amplicon sequencing method. Of the 95 *A. thaliana* transformants expressing mutated *AlSRKb* that contained non-synonymous mutation(s) (and synonymous mutation(s)), 41 showed the SI phenotype and 53 showed the SC phenotype (Table 1; Supplemental Data Set 1). We found that 121 transformants did not synthesize full-length AlSRKb protein; this was due to a mutation in the start codon, a frameshift mutation, a nonsense mutation, or the lack of the *S* domain coding region of *AlSRKb* (Table 1, Supplemental Data Set 1). *AlSRKb* sequences containing only synonymous mutation(s) were found in 17 transformants, and 35 transformants did not contain any mutations in their *AlSRKb* sequences (Table 1; Supplemental Data Set 1). Amplicon sequencing of *AlSRKb* revealed that two different nucleotide sequences were present in 30 transformants, meaning that either two different mutated forms of *AlSRKb* or the *AlSRKb* wild-type plus a mutated form had been introduced (Table 1; Supplemental Figure 1; Supplemental Data Set 1).

### Mutated AlSRKb-FLAG tends not to localize at the plasma membrane in SC transformants

We examined the level of protein expression and the subcellular localization of mutated AlSRKb-FLAG in transformants expressing *AlSRKb* containing non-synonymous mutation(s) (and synonymous mutation(s)); transformants expressing *AlSRKb* containing only synonymous mutation(s) or wild-type *AlSRKb* were used as controls (Figure 2; Supplemental Data Sets 1 and 2). The level of expression in a previously generated *A. thaliana* transformant expressing wild-type AlSRKb-FLAG, which shows an intense SI response (Yamamoto et al. 2014), was set to 100 (Figure 2A; *AlSRKb* WT).

**Figure 2.**
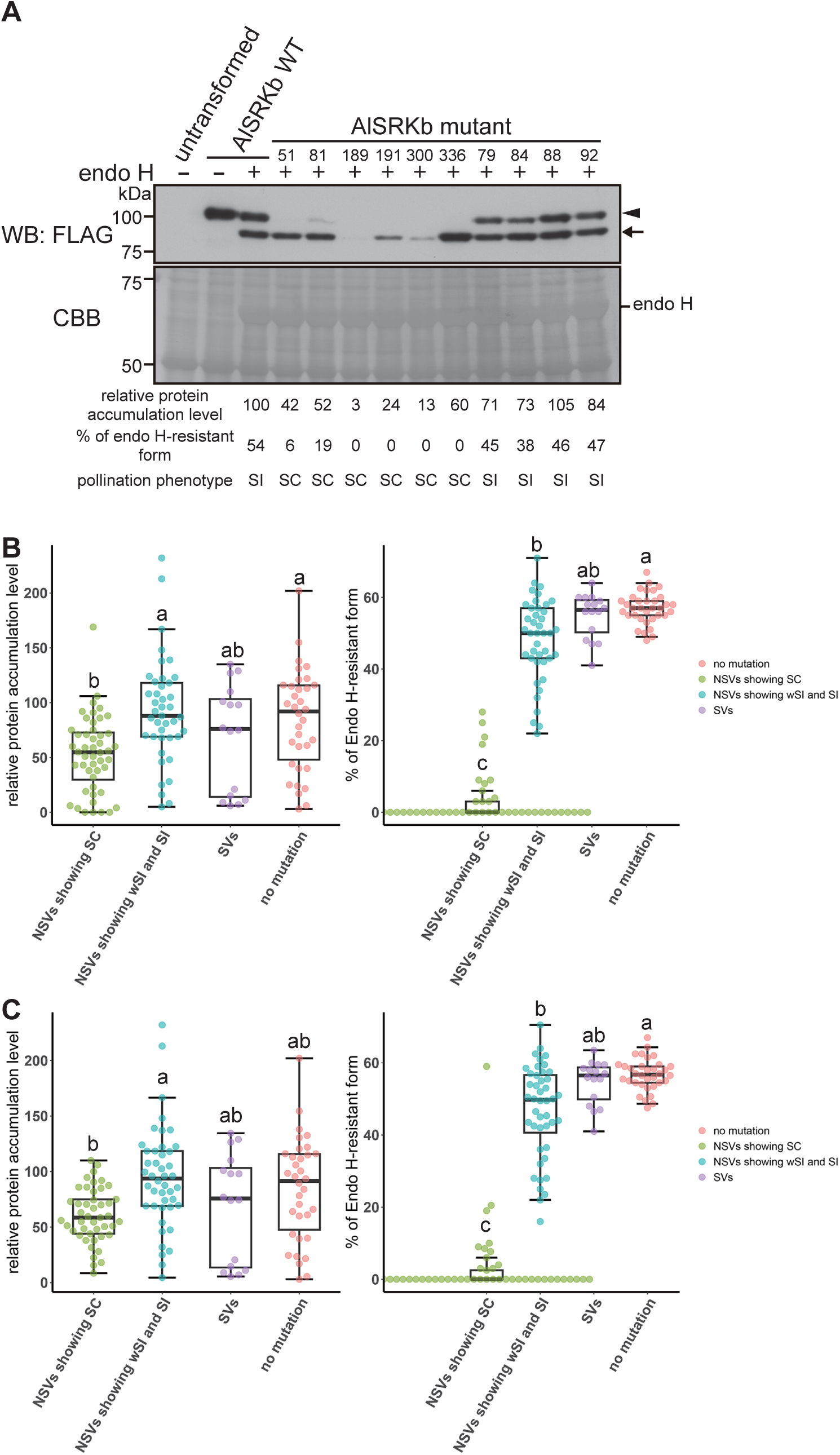
**Determination of expression and plasma membrane localization of AlSRKb mutant proteins in the stigma.** **(A)** Total proteins were extracted from flower buds of untransformed plants (untransformed) and from transformants expressing wild-type AlSRKb-FLAG (AlSRKb WT) or mutated AlSRKb-FLAG (AlSRKb mutant) proteins. The protein samples were incubated with (+) or without (-) endoglycosidase H (Endo-H) at 37 °C for 1 h before being analyzed by western blotting (WB) with anti-FLAG antibody (upper panel). The membrane was then stained with Coomassie brilliant blue (CBB) as a loading control (lower panel). The levels of expression of wild-type and mutated AlSRKb-FLAG proteins, the percentage of protein in the endoglycosidase H-resistant form, and the SI phenotypes are indicated beneath the lanes. The level of expression of wild-type AlSRKb-FLAG protein was set at 100. **(B)** Box plots showing level of protein expression (left) and percentage of the endoglycosidase H-resistant form (right) of AlSRKb-FLAG proteins prepared from stigmas of *A. thaliana* transformants. NSVs showing SC: transformants with a SC phenotype and expressing mutated *AlSRKb-FLAG* genes containing non-synonymous mutation(s), but not mutations resulting in the production of an immature AlSRKb protein; NSVs showing wSI and SI: transformants showing weak SI or SI phenotypes and expressing mutated *AlSRKb-FLAG* genes containing non-synonymous mutation(s); SVs: transformants showing the SI phenotype and expressing mutated *AlSRKb-FLAG* genes containing synonymous mutation(s); no mutation: transformants showing the SI phenotype and expressing wild-type *AlSRKb-FLAG*. Data are means of at least two replicates; full details may be found in Supplemental Data Set 2. Different letters indicate a significant difference between groups, as determined using the Steel-Dwass test (*P* < 0.05). **(C)** The level of protein expression (left) and the percentage of the endoglycosidase H-resistant form (right) of mutated AlSRKb-FLAG proteins prepared from the stigmas of individually constructed *A. thaliana* transformants (Supplemental Data Set 3). These results replicated those shown in Figure 2B; the complete data set may be found in Supplemental Data Set 4. Different letters indicate a significant difference between groups, as determined using the Steel-Dwass test (*P* < 0.05).

The levels of mutated AlSRKb-FLAG in the transformants expressing *AlSRKb* containing non-synonymous mutation(s) showing SC (NSVs showing SC) and SI (NSVs showing SI) were between 0 and 169, and 5 and 232, respectively (Figure 2B left-hand panel; Supplemental Data Set 2). Since some SC transformants accumulated little or no mutated AlSRKb-FLAG (Figure 2B; Supplemental Data Set 2), the SC phenotype of those transformants may result from dysexpression of the mutated protein. To examine this possibility, we constructed individual *A. thaliana* transformants expressing the forms of the mutated *AlSRKb-FLAG* genes from the original transformants #9, #36, #186, #253, #15, #77, and #189, and examined their SI phenotypes using pollination assays with AlSCRb-expressing pollen. Numerous AlSCRb pollen tubes were observed on the stigmas of the newly constructed transformants expressing the mutated *AlSRKb-FLAG* genes #9, #36, #253, #15, and #189 (Supplemental Figure 2A; Supplemental Data Set 3). Furthermore, the levels of accumulation of mutated AlSRKb-FLAG in the novel transformants were detected (Supplemental Figure 2B; Supplemental Data Set 3). These results indicated that the mutated *AlSRKb-FLAG* genes #9, #36, #253, #15, and #189 did not function in SI. By contrast, no or few pollen tubes were observed on the stigmas of the nine transformants expressing the mutated *AlSRKb-FLAG* gene #186 or the four transformants expressing the mutated *AlSRKb-FLAG* gene #77 (Supplemental Figure 2A; Supplemental Data Set 3), suggesting that these two mutated genes conferred SI on *A. thaliana*. We also constructed novel *A. thaliana* transformants expressing the mutated form of *AlSRKb-FLAG* found in transformant #106 (and #109), which showed an SI response and a low level of protein accumulation (Supplemental Data Set 1). In contrast to the original #106 transformant, which contained mutated *ALSRKb-FLAG* produced using error-prone PCR, all the newly constructed transformants expressing mutated *AlSRKb-FLAG* gene #106 showed the SC phenotype (Supplemental Figure 2A; Supplemental Data Set 3).

To determine whether mutated AlSRKb-FLAG proteins were transported to the plasma membrane, we treated proteins extracted from floral buds of *A. thaliana* transformants with endoglycosidase-H (Endo-H). The glycans attached to glycoproteins in the endoplasmic reticulum (ER) are the high mannose-type *N*-glycans in their Endo-H sensitive form. They are modified into the Endo-H resistant form in the Golgi apparatus. A previous study indicated that the result of Endo-H treatment of AlSRKb matches that obtained by subcellular localization analysis of membrane fraction and microscopic observation (Yamamoto et al. 2014). Treatment with Endo-H revealed that Endo-H resistant forms of AlSRKb were either undetected or slightly detected in almost all transformants expressing mutated *AlSRKb* containing non-synonymous mutation(s) and showing an SC phenotype (NSVs showing SC) (Figure 2A,B; Supplemental Data Set 1 and 2), suggesting that mutated AlSRKb was not localized at the plasma membrane in almost all transformants showing defects in SI.

The percentages of the Endo-H resistant form in four transformants, #69, #81, #224, and #227, that showed SC were 25%, 19%, 28%, and 21%, respectively. As this was higher than in other transformants showing SC, we constructed novel *A. thaliana* transformants containing the mutated forms of *AlSRKb-FLAG* from the original transformants #69, #81, #224, and #227 (Supplemental Data Set 3) and examined their SI phenotype using pollination assays. More than 30 pollen tubes were observed germinating from pollen expressing AlSCRb on the stigmas of novel transformants harboring mutated *AlSRKb-FLAG* genes #69 and #227 (Supplemental Figure 2A; Supplemental Data Set 3), indicating that the mutations #69 (K102R, F246I) and #227 (F246L, E287V) produced defective proteins. On the other hand, three of the novel *A. thaliana* transformants expressing mutated *AlSRKb-FLAG* #81 and two transformants expressing #224 showed SI (Supplemental Figure 2A, Supplemental Data Set 3), indicating that mutated AlSRKb proteins #81 (S222G) and #224 (a combination of M313V and T346A) could not abolish SI activity. The SI phenotypes, levels of expression of mutated AlSRKb-FLAG protein, and percentage of the Endo-H-resistant form of the transformants produced using error-prone PCR were replicated in the novel individually generated transformants harboring mutated AlSRKb-FLAG proteins (Figure 2C; Supplemental Data Set 4). In conclusion, our results indicated that *A. thaliana* transformants with a plasma membrane localization rate of less than 10% showed the SC phenotype, suggesting that the threshold for showing the SC phenotype lies between approximately 10% and 20% (Figure 2C; Supplemental Data Set 4).

### The SI defect is caused by 18 mutations in AlSRKb

We next sought to identify the mutations that caused the defect in SI. As 14 of the *A. thaliana* transformants showing SC contained one non-synonymous mutation, we wondered whether these mutations caused the defect in SI (Supplemental Data Sets 4 and 6). The programs SignalP-6.0 (Teufel et al. 2022) and DeepTMHMM (Hallgren et al. 2022) both predicted that 16 mutations found in the signal peptide region would not change signal peptide function (Supplemental Data Set 5), implying that these mutations were not responsible for the changes in AlSRKb function. The transformants #15, #39, and #320 each contained two mutations; as one of these occurred in the signal peptide region, the other mutation was likely to cause the SI defect in these transformants (Supplemental Data Sets 4 and 6). Transformant #90 contained the mutations F53L and E252G. As the E252G mutation was also observed in transformant #24, which showed SI, the F53L mutation may have disrupted AlSRKb function (Supplemental Data Sets 4 and 6). We found that, in total, 18 mutations in AlSRKb led to a defect in SI activity. By contrast, we observed 52 mutations in AlSRKb in transformants that showed an SI or weak SI response (Supplemental Data Sets 4 and 6), which suggested that these mutations were not responsible for the defect in SI activity.

### Characteristics of mutations causing the defect in SI

To explore the properties of the amino acid residues in AlSRKb that are required for the SI response, we compiled the following information on the mutations isolated in this study (Supplemental Data Set 6): “Conservation rate (%) of amino acid residues observed in wild-type AlSRKb and mutated AlSRKb proteins across 49 *S* haplotypes” (Supplemental Figure 3); “BLOSUM62 score (Henikoff and Henikoff 1992) and Grantham’s distance (Grantham 1974) of the mutations found in AlSRKb mutants”, which show how point mutations change amino acid properties; “Relative accessible surface area (RASA) of amino acid residues in an AlSRKb structural model produced by ColabFold (Mirdita et al. 2022)”, “DeepDDG score” and “STRUM score”, which show how point mutations alter the stability of an AlSRKb structural model, as predicted by, respectively, the DeepDDG (Cao et al. 2019) and STRUM (Quan et al. 2016) programs; “Hydrophobicity (Monera et al. 1995) of the amino acid residues observed in wild-type AlSRKb and mutated AlSRKb proteins”, “pI (Lide 1991) of the amino acid residues observed in wild-type AlSRKb and mutated AlSRKb proteins”, “Predicted amino acid residues in AlSRKb involved in the interaction with AlSCRb or in AlSRKb homodimerization”, “Amino acid residues in AlSRKb forming disulfide bridges”, and “Amino acid residues required for *N*-glycosylation”. Scatter plots revealed that the values for “Conservation rate (%) of amino acid residues observed in wild-type AlSRKb and mutated AlSRKb proteins across *S* haplotypes”, “BLOSUM62 score”, “Grantham’s distance”, “RASA”, “DeepDDG score”, and “STRUM score” differed between transformants showing the SC and SI phenotypes (Supplemental Figure 4). This analysis identified four features of AlSRKb mutations causing defects in SI: first, the mutation occurred in amino acid residues that were highly conserved across *S* haplotypes; second, the mutation caused significant changes in the amino acid properties; third, the mutation was located on the interior of the AlSRKb protein; and fourth, the mutation had a destabilizing effect on AlSRKb structure.

Next, we used the parameters discussed above in a principal component analysis (PCA) (Figure 3A; Supplemental Data Set 6). The result of PCA analysis, which showed in the 24.7 % (PC1) and 13.1% (PC2) variance with two-component analysis, revealed that mutations causing SC were usually separated from those causing SI, although some mutations from each group were overlapped (Figure 3A). This separation may result from the parameters “AlSRKb structure model stability changes by a point mutation (DeepDDG and STRUM scores)”, “Conservation rate (%) of the amino acid residues observed in wild-type AlSRKb across *S* haplotypes”, and “the change in amino acid properties caused by a point mutation (BLOSUM62 score and Grantham’s distance)”.

**Figure 3.**
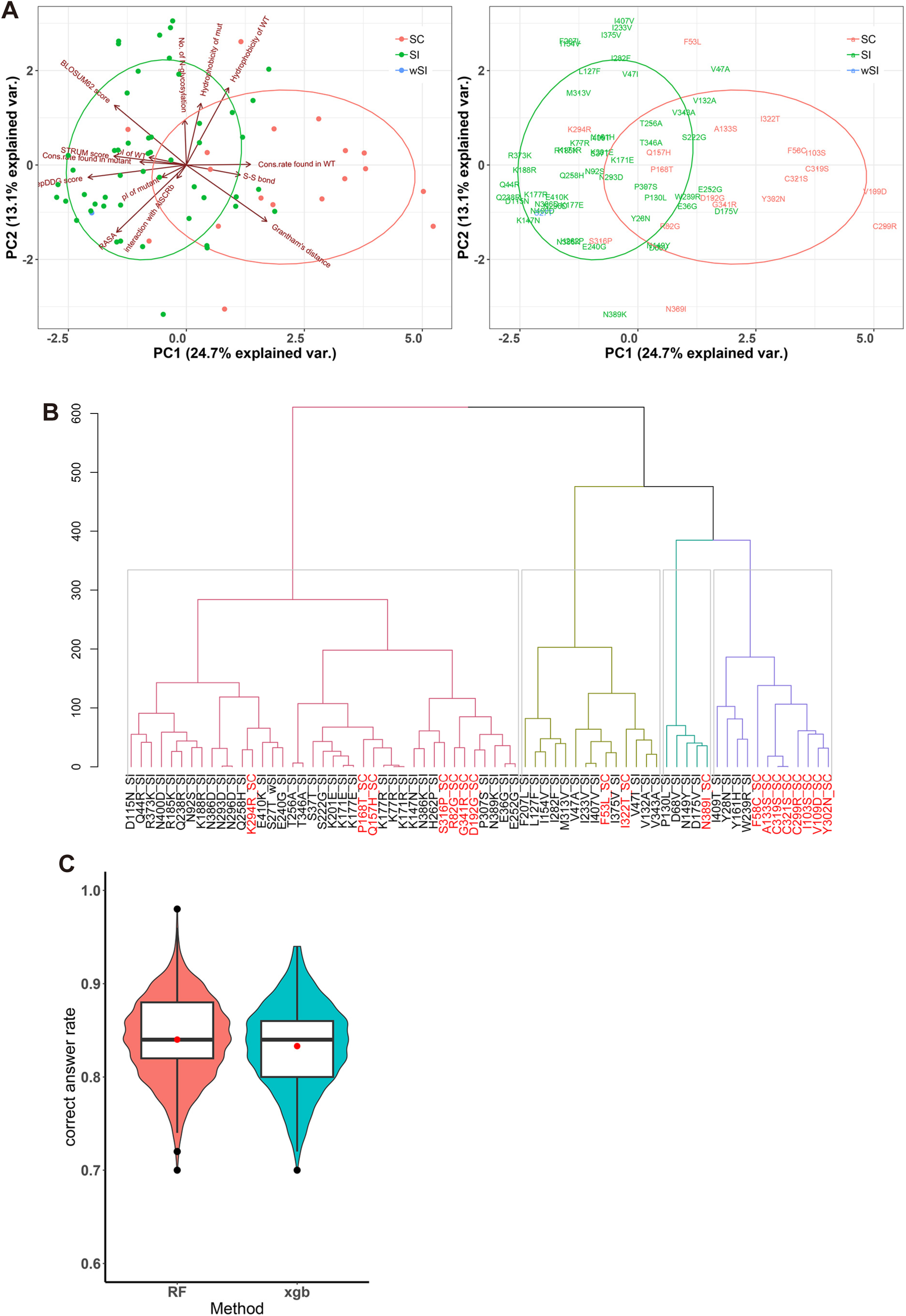
**Prediction of phenotypes associated with different mutations in AlSRKb proteins using machine learning methods.** **(A)** PCA analysis of AlSRKb mutations isolated in this study. Mutations causing the defect in SI are shown in magenta; mutations that did not cause the defect in SI are shown in green. The mutation shown in blue (S27T) was found in *A. thaliana* transformants showing a weak SI phenotype. The specific mutations associated with each dot (left-hand panel) are shown in the right-hand panel. All the data used for PCA analysis are in Supplemental Data Set 6. **(B)** Cluster analysis of the AlSRKb mutations used in the PCA analysis. **(C)** Violin plots showed the correct answer rates produced by the RandomForest (RF) and Extreme Gradient Boosting (xgb) methods. Seventy percent of the mutations listed in Supplemental Data Set 8 were used as training data and the other 30% as test data. The SI phenotypes for the test data were predicted using the RF and Extreme Gradient Boosting (xgb) methods, and compared with the phenotype determined by pollination assay. One thousand bootstrap replicates were performed. Red dots indicate the mean values.

We also performed a cluster analysis using the same parameters as for the PCA analysis. This showed that eight of the 18 mutations that caused the defect in SI clustered together, while the others formed branches with mutations observed in transformants showing the SI phenotype (Figure 3B).

### Prediction of mutations causing the defect in SI

In total, 58 mutations were found in *A. thaliana* transformants expressing mutated *AlSRKb-FLAG* that showed the SC phenotype and had two or more non-synonymous mutations. As it was unknown which of the mutation(s) were responsible for the defect in SI, we used machine learning methods to predict which mutations were likely to underlie this response. For this purpose, in addition to the results of this study, we used the SI phenotypes of SRKs mutants reported previously (Boggs et al. 2009, Yamamoto et al. 2014). Yamamoto et al. 2014 mainly focused on the *N*-glycosylation sites in AlSRKb (Supplemental Data Set 7). Boggs et al. 2009 induced mutations in amino acid residues of chimeric SRK proteins consisting of either *A. lyrata* SRKa and *Capsella grandiflora* SRK7 (hereafter, eSRKa(7)a), or *A. lyrata* SRK16 and *A. lyrata* SRK25 (hereafter, eSRK16(25)16). They focused on the hypervariable (hv) regions of the chimeric proteins. These data from Boggs et al. 2009 provide a useful supplement to the present study, which does not provide much information about mutations in the hv regions of SRK. We therefore compiled the parameters of eSRKa(7)a and eSRK16(25)16 (Supplemental Data Set 7).

To check the rate of correct answers offered by our prediction methods, we divided the mutations whose SI phenotypes were known (Supplemental Data Set 8) at random into two parts in a 7:3 ratio, using the larger part of the mutations (70%) as the training data set and the rest (30%) as the test data set, and used the training data to predict the SI phenotypes of the mutations in the test data set; these predictions were repeated 1,000 times. The predicted SI phenotypes were compared with the phenotypes determined by experiment to obtain the correct answer rate (Figure 3C). The mean values of the correct answer rate were 84.0% (RandomForest method) and 83.3% (Extreme Gradient Boosting method) (Figure 3C).

We next predicted the SI phenotypes of the 58 mutations whose actual SI phenotype could not be determined by experiment because multiple mutations occurred within the same transformant (Supplemental Data Set 9). These predictions were repeated 1,000 times to determine the frequencies of the predicted SI and SC phenotypes (Supplemental Data Set 9). A phenotype predicted more than 800 times (80%) was considered the consensus predicted phenotype (Supplemental Data Set 9).

The two methods predicted the same consensus phenotype for 55 of the 58 mutations; 36 mutations were predicted to produce the SI phenotype and 19 mutations the SC phenotype (Supplemental Data Set 9). The mutation S158G was predicted as SC by the Extreme Gradient Boosting program, but this was not the consensus phenotype predicted by the RandomForest program (Supplemental Data Set 9); two mutations (F246I and A394G) were, however, predicted as the SC phenotype by the RandomForest program, but not by the Extreme Gradient Boosting program (Supplemental Data Set 9).

To evaluate these predictions, we constructed novel *A. thaliana* transformants expressing the *AlSRKb*(*N89D*), *AlSRKb*(*M170T*), and *AlSRKb*(*G173D*) mutations; all were predicted to show the SC phenotype (Supplemental Data Set 9). During pollination assays, more than 30 pollen tubes from AlSCRb-expressing pollen were observed on the stigmas of all the novel transformants (Figure 4A, Table 3); this result is consistent with the predicted phenotypes. Endo-H treatment revealed that the AlSRKb(N89D), AlSRKb(M170T), and AlSRKb(G173D) mutant proteins were all Endo-H-sensitive forms (Figure 4B). Taken together, these results indicated that the mutations N89D, M170T, and G173D caused the defects in SI activity and plasma membrane localization of AlSRKb.

**Figure 4.**
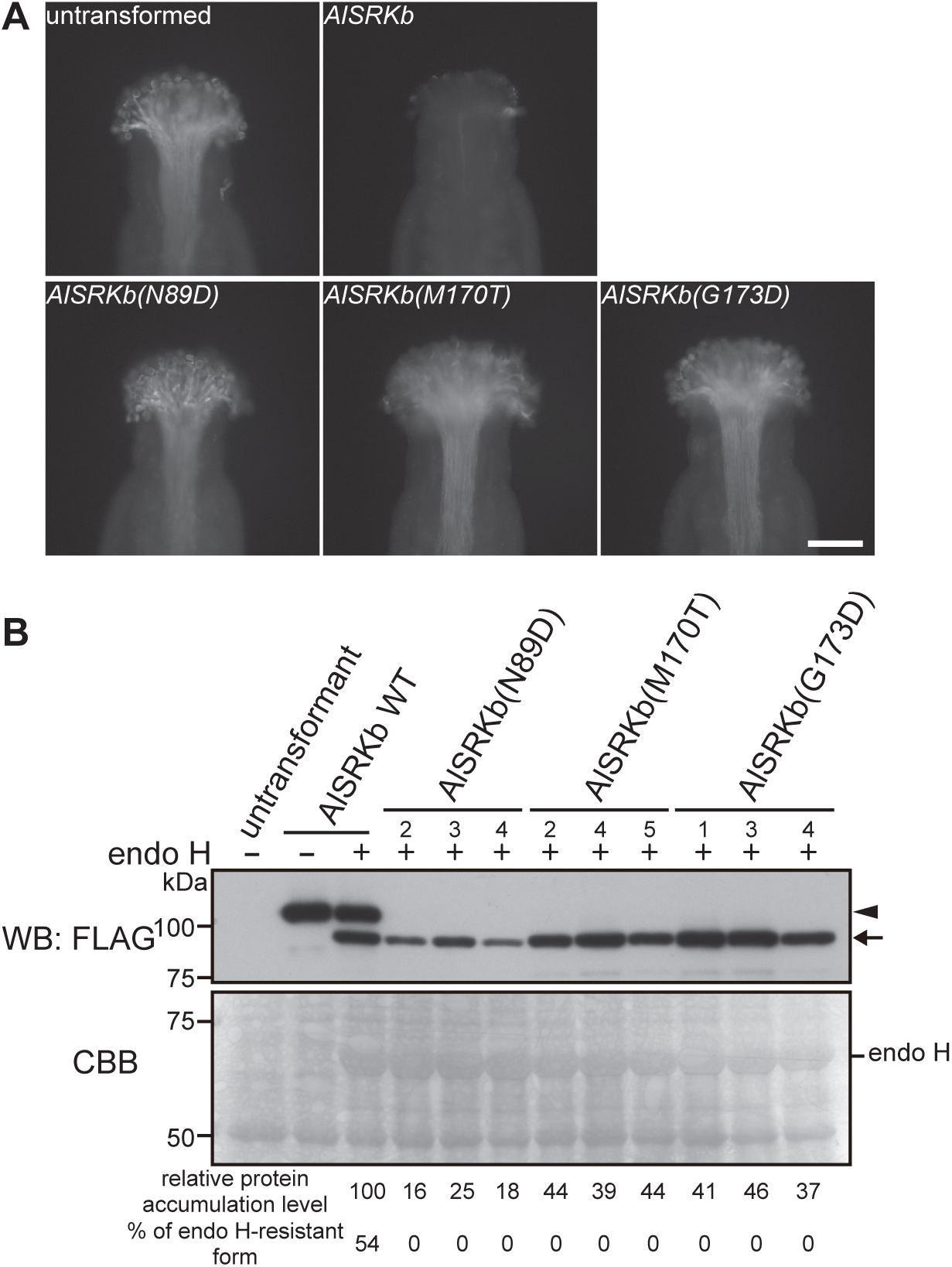
**Evaluation of the effects of the N89D, M170T, and G173D mutations on SI activity.** **(A)** Microscopic observations of pollinated stigmas from *A. thaliana* plants showing AlSCRb-expressing pollen stained with aniline blue. Untransformed and *AlSRKb* are as described in Figure 1. *AlSRKb*(*N89D*), *AlSRKb*(*M170T*), and *AlSRKb*(*G173D*) indicate *A. thaliana* transformants expressing *AlSCRb* + *AlSRKb-FLAG*(*N89D*), (*M170T*), and (*G173D*), respectively. Scale bar = 100 μm. **(B)** Level of protein expression and the percentage of the endoglycosidase H-resistant form of the AlSRKb(N89D), AlSRKb(M170T), and AlSRKb(G173D) proteins; experiments were performed as described in Figure 2A.

**Table 2.**
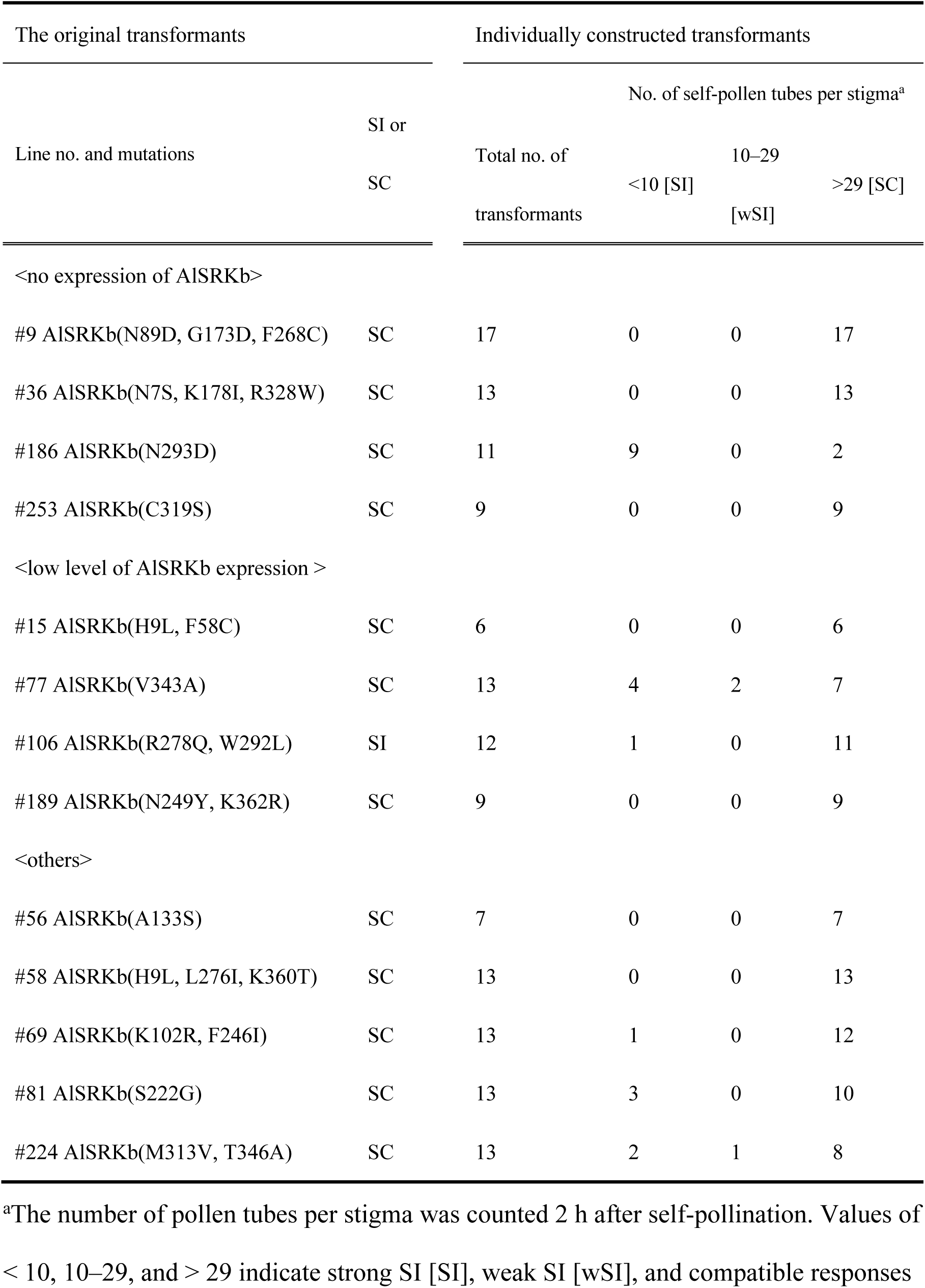
SI phenotypes of individually constructed *A. thaliana* transformants expressing mutated *AlSRKb*.

**Table 3.** SI phenotypes of *A. thaliana* transformants expressing *AlSRKb(N89D)*, *AlSRKb(M170T)*, and *AlSRKb(G173D)*

| Transgene | Total no. of transformants | No. of self-pollen tubes per stigma<sup>a</sup> |  |  |
| --- | --- | --- | --- | --- |
|  |  | <10 [SI] | 10–29 [wSI] | >29 [SC] |
| *AlSRKb(N89D) FLAG* | 10 | 0 | 0 | 10 |
| *AlSRKb(M170T) FLAG* | 11 | 0 | 0 | 11 |
| *AlSRKb(G173D) FLAG* | 9 | 0 | 0 | 9 |

<sup>a</sup>The number of pollen tubes per stigma was counted 2 h after self-pollination. Values of < 10, 10–29, and > 29 indicate strong SI [SI], weak SI [wSI], and compatible responses [SC], respectively.

## DISCUSSION

This study revealed that almost all *A. thaliana* transformants with a SC phenotype showed defective localization of mutated AlSRKb-FLAG to the plasma membrane. As newly synthesized proteins usually fold properly before leaving the ER (Braakman and Hebert 2013), these mutated AlSRKb-FLAG proteins were likely to have disruptions in their protein folding. Amino acid residues with hydrophobic side chains tend to cluster in the center of protein molecules to avoid exposure to the aqueous environment (Chothia 1974); therefore, protein structure is often destabilized by mutations in amino acid residues located in the interior of proteins. In this study, many of the mutated AlSRKb proteins in *A. thaliana* transformants showing SC were found to harbor mutations in amino acids in the interior of the protein structural model. We have, however, previously reported that mutations in BrSRK9, including BrSRK9(V211E) and BrSRK9(P294M), whose expression does not confer SI activity in *A. thaliana*, at amino acid residues involved in the BrSRK9-BrSCR9 interaction does not affect localization to the plasma membrane (Yamamoto et al. 2022). This implies that mutations causing defects in SI and in the interaction with SCR will not also affect SRK localization. Our current study suggested, however, that mutations in AlSRKb causing defects in SI were more likely to disrupt AlSRKb biosynthesis than those that resulted in a loss of AlSRKb function, such as dimerization and interaction with AlSCRb.

The RandomForest and Extreme Gradient Boosting methods predicted the phenotypes caused by mutations with 84.0% and 83.3% accuracy, respectively. The predicted phenotypes (SC) of transformants expressing *AlSRKb(N89D)*, *AlSRKb(M170T)*, and *AlSRKb(G173D)* were confirmed experimentally using pollination assays. The causative mutation(s) in 27 transformants showing the SC phenotype were unknown because the AlSRKb proteins in these plants contained multiple mutations (Supplemental Data Set 10). Candidates for causative mutations in 18 of the 27 transformants were predicted: 19 mutations were predicted to produce the SC phenotype by both the RandomForest and Extreme Gradient Boosting methods, and one mutation was classified as producing SC only by the Extreme Gradient Boosting method (Supplemental Data Set 10). No candidate causative mutations were predicted for the other nine transformants (#36, 58, 68, 69, 85, 127, 197, 199, and 227) (Supplemental Data Set 10). Supplemental Data Set 11 shows 10 examples of the phenotypes predicted for specific mutations during the analysis of the data shown in Figure 3C. For 16 mutations (V47A, R82G, Q157H, P168T, K294R, S316P, N389I, and T391A in AlSRKb; L218V, D299E, Y300T, and I355T in eSRKa(7)a; K221F, Q293E, S300F, and H303K in eSRK16(25)16), all the predicted phenotypes were answered incorrectly in multiple tests (Supplemental Data Set 11). In 14 out of 16 mutations known to cause the defect in SI, the phenotype was incorrectly predicted as SI, meaning that our predictive methods missed mutations causing SC. The hv regions contained seven mutations; some amino acid residues in these regions are involved in SRK homodimerization and the interaction with SCR (Ma et al. 2016, Murase et al. 2020, Sawa et al. 2023). These amino acid residues are located at the surface of AlSRKb and some appear to play an important role in SI activity (Boggs et al. 2008, Yamamoto et al. 2022). Although our results suggested that mutations in amino acid residues that were highly conserved across *S* haplotypes tended to cause the defect in SI, amino acid residues within the hv regions are generally not conserved across *S* haplotypes. Our PCA analysis showed that the mutations K294R and S316P, which caused the defect in SI, were located in the group of transformants showing the SI phenotype, and the V47A, R82G, Q157H, and P168T mutations were located in the overlap region. The mutations N389I and T391A occurred in the *N*-glycosylation site. Based on these results, predicting the effect of mutations that show characteristics other than those generally associated with the defect in SI, as described in the Results section, becomes more difficult. It is therefore necessary to improve the accuracy with which we can predict SI phenotypes associated with mutations showing different characteristics.

The present study suggested that the SI phenotype associated with mutations in in the amino acid residues located outside the hv regions of AlSRKb could be predicted with high accuracy. As amino acid residues outside the hv regions are relatively highly conserved across *S* haplotypes, these results improve our ability to predict which amino acid residues are essential for SI activity, not only of AlSRKb but also of other SRK proteins. In addition, the results of this study will inform future searches for functionally important SNVs and mutations in both *SRK* genes and other genes, and thus contribute to genetic analysis by aiding the identification of causative genes underlying important genetic traits.

## METHODS

### Plant material and growth conditions

*Arabidopsis thaliana* ecotype C24, obtained from the Arabidopsis Biological Resource Center (ABRC; https://abrc.osu.edu/), was used as the wild-type plant in this study. The *AlSRKb-FLAG*+*AlSCRb A. thaliana* transformants were constructed as described in Yamamoto et al. (2014). Seeds were surface-sterilized with 20% bleach and sown on Murashige and Skoog medium (FUJIFILM Wako, Osaka, Japan) containing 0.7% (w/v) agar and 1% (w/v) sucrose. All *A. thaliana* plants were grown at a constant temperature of 23 °C under a long-day photoperiod (16 h light/8 h dark; light intensity 100 mmol photons m^-2^ sec^-1^). The relative humidity was not controlled.

### Construction of a plasmid library containing mutated *AlSRKb-FLAG* genes with randomly introduced mutations and a population of *A. thaliana* transformants expressing mutated *AlSRKb-FLAG* genes

A schematic diagram of the construction of the population of *A. thaliana* transformants expressing mutated *AlSRKb-FLAG* is shown in Figure 1A. The construction of the *AtS1_pro_*:*AlSRKb FLAG*+*AlSCRb* transgene, which is derived from the pCAMBIA1300 plant transformation plasmid (GenBank accession number AF234296), has been described previously (Yamamoto et al., 2014). Comparison of the *AlSRKb* nucleotide sequence deposited in Genbank (GenBank accession number AB052756) with the *AlSRKb* sequence in the *AtS1_pro_*:*AlSRKb FLAG*+*AlSCRb* transgene revealed that, at position 1,100, C was replaced with T, thus the wild-type AlSRKb protein used in this study contained an S367L mutation not present in the protein sequence deposited in Genbank (GenBank accession number BAB40987).

The mutations in the DNA sequence of the *AlSRKb* receptor domain coding region were generated by the error-prone PCR method using homemade *Taq* (Pluthero, 1993), a reaction buffer (10 mM TrisHCl pH8.3, 50 mM KCl, 7 mM MgCl_2_, 50 mM MnCl_2_, 0.8 mM dNTPs), the AtS1(-28)/AlSRKb(1260)R primer pair (Supplemental Data Set 13), and the *AtS1_pro_*:*AlSRKb FLAG*+*AlSCRb* transgene as a template. PCR conditions were 94 °C for 2 min, followed by 20 cycles of 94 °C for 30 sec, 50 °C for 30 sec, and 72 °C for 3 min.

The amplified DNA fragment was introduced into the *Kpn*I/*Sac*I sites of the *AtS1_pro_*:*AlSRKb FLAG*+*AlSCRb* transgene using the LigaFast Rapid DNA Ligation System (Promega, Fitchburg, WI, USA). The ligation products were transformed into *Escherichia coli* DH5a competent cells. Plasmid DNA was extracted from approximately 1,000 colonies, which were cultivated in the same flask. The plasmids were used to transform *Agrobacterium tumefaciens* strain GV3101 (Koncz and Schell, 1986) and approximately 7,000 colonies were cultivated in the same flask to construct an Agrobacterium library. The floral dip method (Clough and Bent, 1998) was used to transform *A. thaliana* C24 ecotype with the Agrobacterium library. Transformants were selected on Murashige and Skoog medium containing 50 mg/mL hygromycin.

### Determination of mutations in *AlSRKb-FLAG* genes expressed in *A. thaliana* transformants

Genomic DNA was prepared from leaves of *A. thaliana* transformants expressing mutated *AlSRKb-FLAG* using the cetyltrimethyl ammonium bromide (CTAB) method (Doyle and Doyle, 1987). PCR was used to amplify four DNA fragments from each mutated *AlSRKb-FLAG* gene using KOD-Plus-Neo (TOYOBO, Osaka, Japan), the appropriate primers, and the genomic DNA template under the following amplification conditions: 94 °C for 2 min, followed by 35 cycles of 94 °C for 15 sec, 58 °C for 30 sec, and 68 °C for 30 sec. The primer pair SRKb-1stF/SRKb-1stR was used to amplify the region of the *AlSRKb* coding sequence (CDS) between base pairs (bp)-28 and 330; the primer pair SRKb-2ndF/SRKb-2ndR to amplify the CDS region between bp 274 and 621; the primer pair SRKb-3rdF/SRKb-3rdR to amplify the CDS region between bp 579 to 934; and the primer pair SRKb-4thF/SRKb-4thR to amplify the CDS region between bp 874 and 1,283 (Supplemental Data Set 13). Index sequences, as described previously (Shirasawa et al 2016), were attached to the amplified products by PCR using KOD-Plus-Neo (TOYOBO) under the same conditions as the first round of PCR. The resulting amplicons were pooled and purified by NucleoSpin Gel and PCR Clean-up (TAKARA Bio, Shiga, Japan).

The purified fragments were sequenced on the MiSeq platform (Illumina, San Diego, CA) to generate 300 bp paired-end reads. Low-quality and adapter sequences were removed using fastp (Shifu et al. 2018) and the filtered reads were mapped onto the *AlSRKb* CDS sequence (AB052756) using BWA-MEM (Li, 2013). The resulting sequence alignment/map (SAM) format files were converted to binary sequence alignment/map format files and visualized using the Integrative Genomics Viewer (IGV) (James et al. 2017). The mutations were manually searched. All the amplicon sequencing data described here have been deposited in the DDBJ BioProject database under BioProject accession number PRJDB17639.

### Plasmid construction and generation of transgenic *A. thaliana* plants

Genomic DNA was extracted from the leaves of *A. thaliana* transformants #9, 15, 36, 69, 77, 81, 106, 186, 189, 224, 227, and 253. Fragments of the mutated *AlSRKb* genes from the transformants were amplified by PCR using the primer pair AtS1(-28)/AlSRKb(1260)R (Supplemental Data Set 13). The amplified fragments were introduced into the *Kpn*I/*Sac*I sites of *AtS1_pro_*:*AlSRKb FLAG*+*AlSCRb* using the LigaFast Rapid DNA Ligation System (Promega). The mutations N89D, M170T, and G173D were introduced into the *AlSRKb* sequence by recombinant PCR-mediated site-directed mutagenesis using *AtS1_pro_*:*AlSRKb FLAG*+*AlSCRb* as a template and the appropriate sequence-specific primer pair: (AlSRKb(N89D)F/AlSRKb(N89D)R for *AlSRKb(N89D)*; (AlSRKb(M170T)F/AlSRKb(M170T)R for *AlSRKb(M170T)*; and AlSRKb(G173D)F/AlSRKb(G173D)R for *AlSRKb(G173D)* (Supplemental Data Set 13).

The resulting plasmids were sequenced at Eurofins Genomics KK (Tokyo, Japan) to confirm the absence of PCR-generated polymorphisms. The sequence-verified plasmids were used to transform *Agrobacterium tumefaciens* strain GV3101 (Koncz & Schell, 1986) and introduced into wild-type C24 plants by the floral dip method (Clough and Bent, 1998). Transformants were selected on Murashige and Skoog medium containing 50 mg/mL hygromycin.

### Pollination assays

*A. thaliana* flowers were emasculated just before flowering, which corresponds with flowering stage 13 (Smyth et al., 1990), and placed on a 0.5% agar plate. Stigmas were manually pollinated under a stereomicroscope with pollen grains from open flowers. The pollinated stigmas were incubated at 23 °C under continuous light for 2 h before fixation in ethanol:acetic acid (3:1) for 10 min at 55 °C. The fixed stigmas were treated with 8 M NaOH at room temperature for 15 min and then stained with decolorized aniline blue. The stained stigmas were observed by fluorescence microscopy, as previously described (Kho and Bear, 1968). At least three stigmas were used in each pollination assay.

The degree of SI activity was classified according to the number of pollen tubes per pollinated stigma as follows: strong SI < 10 pollen tubes; weak SI 10–29 pollen tubes; compatible > 29 pollen tubes. Images of pollinated stigmas were captured using an Axioskop microscope with an AxioCam ERc 5 s camera (Carl Zeiss, Oberkochen, Germany).

### Western blot analysis

Samples of 10 *A. thaliana* buds in positions-1 (bud immediately previous to the flower) and-2 (the second bud before the flower) were collected and homogenized in extraction buffer (100 mM Tris–HCl, pH 8.0, 2% [w/v] SDS, 5.7 mM b-mercaptoethanol, and 1 mM phenylmethylsulfonyl fluoride). Extracted proteins were precipitated using the trichloroacetic acid/acetone method and dissolved in SDS-PAGE sample buffer (Laemmli, 1970). The protein samples were treated with Endo Hf (New England BioLabs, Ipswich, MA, USA) at 37°C for 1 h. The proteins were separated by 7.5% SDS-PAGE and transferred to an Immobilon-P membrane (Millipore, Billerica, MA, USA), as previously described (Towbin et al., 1979). Proteins were detected using anti-FLAG (Sigma-Aldrich, ST. Louis, MO, USA) and HRP-labeled anti-mouse IgG (Cytiva, Tokyo, Japan) at 1:5,000 dilution. Immunodetection was performed using Pierce ECL Plus Western Blotting Substrate (ThermoFisher Scientific, Waltham, MA, USA) and exposure to X-ray film. The detected proteins were quantified using ImageJ software (https://imagej.nih.gov/ij/). The membranes were subsequently stained with Coomassie Brilliant Blue R250.

### Data analysis

The Steel-Dwass test was performed using the python package scikit-posthocs (Terpilowski, M., 2019) for multiple comparisons of expression level and the ratio of the endoglycosidase H-resistant form of mutated AlSRKb proteins in Figure 2B,C. The box plots and scatter plots were drawn using R package ggplot2 (Wickham, H. 2016). Principal component analysis (PCA) was performed using the function prcomp in R and visualized using the ggbiplot package. The cluster analysis was performed in the R stats package using the function hclust with the ward.D2 method.

Structural models of the *S*-domains of the AlSRKb, eSRKa(7)a, eSRK16(25)16, AlSCRb, *Capsella grandiflora* SCR7 (CgSCR7), and AlSCR25 proteins were generated using ColabFold (Mirdita, M. et al. 2022). The solvent accessible surface area of the predicted SRK models was calculated using the DSSP program (Kabsch W. and Sander C. 1983, Wouter et al. 2015). The relative accessible surface area (RASA) for each amino acid residue was determined by dividing the calculated solvent accessible surface area by a theoretical solvent accessible surface area, obtained from Tien et al. 2013, for each amino acid residue. DeepDDG scores and STRUM scores were calculated for the predicted SRK structural models using the DeepDDG (Cao et al. 2019) and STRUM (Quan et al. 2016) programs. To determine which amino acid residues in SRK proteins were involved in the interaction with SCR or SRK homodimerization, the two *S*-domains of the SRK models and the two SCR models were superimposed on the BrSRK9-BrSCR9 structure (PDB id: 5GYY), and amino acid residues of one SRK model within 5 Å of the other SRK model and the two SCR models were selected using UCSF ChimeraX (Meng et al. 2023). BLOSUM62 scores and Grantham’s distance were obtained from Henikoff S and Henikoff JG (1992) and Grantham R (1974), respectively.

Signal sequences of mutated AlSRKb proteins were predicted using the SignalP-6.0 (Teufel et al. 2022) and DeepTMHMM (Hallgren et al. 2022) programs. Alignments of amino acid sequences of the SRK variants were produced by MEGA11 (Tamura et al. 2021) and visualized using SeaView (Gouy et al. 2009).

### Phenotype prediction

Phenotype (SI or SC) was predicted using the RandomForest (R package RandomForest (Liaw and Wiener, 2002)) and Extreme Gradient Boosting (R package xgboost (Tianqi and Carlos, 2016)) methods. To simplify the prediction, the weak SI phenotype observed in the pollination assays of *A. thaliana* transformants expressing AlSRKb(S27T) (Supplemental Data Set 6) was treated as SI (Supplemental Data Set 8). In addition to the mutations found in the present study, the SI phenotypes of previously reported SRK mutants (Boggs et al. 2009, Yamamoto et al. 2014) were included.

To evaluate the accuracy of the prediction methods, 70% of the mutations, whose effects on the SI phenotype were known from the pollination assays (Supplemental Data Set 8), were used as training data and the other 30% as test data. The SI phenotypes associated with mutations in the test data set were predicted by both methods using the training data. The predicted phenotypes were compared with the actual phenotypes, and the correct answer rates of the predictions after 1,000 bootstrap replicates were visualized using the ggplot2 package (Wickham, H. 2016).

The RandomForest and Extreme Gradient Boosting methods were used to predict the phenotypes of mutations whose actual effects on SI were unknown because they occurred in association with other mutations (Supplemental Data Set 9). Mutations with known effects on phenotype (SI or SC; Supplemental Data Set 8) were used as a training data set with 1,000 replicates. If SI was predicted in > 800 of 1,000 replicates (80%), the predicted phenotype was classified as SI [SI]; if > 80% replicates predicted SC, then the predicted phenotype was classified as SC [SC]; if no clear result was obtained from 1,000 replicates, the predicted phenotype was classified as unknown [?].

## ACKNOWLEDGMENTS

Protein structure graphics and analyses were performed using UCSF ChimeraX, developed by the Resource for Biocomputing, Visualization, and Informatics at the University of California, San Francisco, with support from National Institutes of Health R01-GM129325 and the Office of Cyber Infrastructure and Computational Biology, National Institute of Allergy and Infectious Diseases. This work was supported by the Japan Society for the Promotion of Science KAKENHI (Grant Numbers 20K05979 and 23K05161 to MY), a Cooperative Research Grant for Genome Research for BioResource, NODAI Genome Research Center, Tokyo University of Agriculture (Grant Numbers 2019_B013 to HK), and Research Support Project for Life Science and Drug Discovery (Basis for Supporting Innovative Drug Discovery and Life Science Research (BINDS)) from AMED under Grant Number JP23ama121019.

## AUTHOR CONTRIBUTIONS

MY designed the research; MY, SO, AS, and MS performed the research and analyzed the data. All authors wrote the manuscript.

## Supplemental data

The following materials are available in the online version of this article.

**Supplemental Figure S1.** Examples of mutated AlSRKb DNA sequences.

**Supplemental Figure S2.** Phenotypic analysis of A. thaliana transformants expressing mutated AlSRKb-FLAG.

**Supplemental Figure S3.** Alignment of amino acid sequences of the SRK variants from Arabidopsis lyrata (AlSRKx), A. halleri (AhSRKx), and *Capsella grandiflora* (CgSRKx) used in this study.

**Supplemental Figure S4.** Comparison of the values of mutations found in A. thaliana transformants showing the SC and SI phenotypes.

**Supplemental Data Set 1.** List of genotypes and phenotypes of *Arabidopsis thaliana* transformants expressing mutated *SRKb* from *Arabidopsis lyrata*.

**Supplemental Data Set 2.** Expression levels and percentages of the endoglycosidase H-resistant forms of mutated AlSRKb proteins shown in Figure 2B.

**Supplemental Data Set 3.** Phenotype, level of protein expression, and percentage of endoglycosidase H-resistant form in transformants expressing individual forms of mutated *AlSRKb*.

**Supplemental Data Set 4.** Expression level and percentage of the endoglycosidase H-resistant form of mutated AlSRKb proteins shown in Figure 2C.

**Supplemental Data Set 5.** Predicted signal peptides and their cleavage sites in mutated AlSRKb proteins.

**Supplemental Data Set 6.** List of AlSRKb mutations found in this study.

**Supplemental Data Set 7.** List of previously reported mutations in SRK proteins.

**Supplemental Data Set 8.** List of mutations in the SRK proteins used as a training data set for phenotype predictions.

**Supplemental Data Set 9.** Phenotype predictions for mutated AlSRKb proteins.

**Supplemental Data Set 10.** Phenotype predictions for *A. thaliana* transformants expressing mutated *AlSRKb*.

**Supplemental Data Set 11.** Ten examples of phenotypes predicted for mutated AlSRKb proteins.

**Supplemental Data Set 12.** List of mutated SRK proteins whose phenotypes were predicted incorrectly in Supplemental Data Set 11.

**Supplemental Data Set 13.** List of oligonucleotide primers used in this study.

## Supplemental Figure legends

**Supplemental Figure S1.** Examples of mutated *AlSRKb* DNA sequences. The DNA sequences of mutated *AlSRKb* genes from *A. thaliana* transformants #8, 157, and 315 were determined by amplicon sequencing and the nucleotide region between base pairs 582 and 674 of *AlSRKb* were visualized using the Integrative Genomics Viewer (IGV). The mutation A645G was observed in almost all *AlSRKb* genes in transformant #315 (shown in green). By contrast, the mutations T606C and T637C were observed in approximately 50% of *AlSRKb* genes in transformants #8 and #157 (shown in yellow).

**Supplemental Figure S2.** Phenotypic analysis of *A. thaliana* transformants expressing mutated *AlSRKb-FLAG*.

**(A)** Microscopic observations were performed as described in Figure 1. Note that no or few pollen tubes germinated from AlSCRb-expressing pollen on the stigmas of transformants expressing mutated *AlSRKb-FLAG* genes #186, #77, #81, and #224, indicating that these transformants showed an SI response. Scale bar = 100 μm.

**(B)** Level of protein expression and the percentage of the endoglycosidase H-resistant form of mutated AlSRKb proteins in freshly constructed individual *A. thaliana* transformants expressing specific forms of mutated *AlSRKb-FLAG* genes. Experiments were performed and analyzed as shown in Figure 2A.

**Supplemental Figure S3.** Alignment of amino acid sequences of the SRK variants from *Arabidopsis lyrata* (AlSRKx), *A. halleri* (AhSRKx), and *Capsella grandiflora* (CgSRKx) used in this study.

**Supplemental Figure S4.** Comparison of the values of mutations found in *A. thaliana* transformants showing the SC and SI phenotypes. The values for “Conservation rate (%) of amino acid residues observed in wild-type and mutated AlSRKb proteins across *S* haplotypes”, “Grantham’s distance” and “BLOSUM62 score”, “Relative accessible surface area (RASA) of amino acid residues in the AlSRKb structural model”, “DeepDDG score”, “STRUM score”, “Hydrophobicity of amino acid residues observed in wild-type and mutated AlSRKb proteins”, and “pI of amino acid residues in wild-type and mutated AlSRKb proteins” of mutations present in *A. thaliana* transformants showing the SC and SI phenotypes were compared. Welch’s *t* tests were performed to determine statistically significant differences between groups.

## SUPPORTING INFORMATION

Additional Supporting Information may be found in the Supporting Information section at the end of the online version of the article.

## Notes

### Competing Interest Statement

The authors have declared no competing interest.

