## Supplemental Figures for "Self-incompatibility phenotypes of SRK mutants can be predicted with high accuracy"

Fig S1 Yamamoto et al.

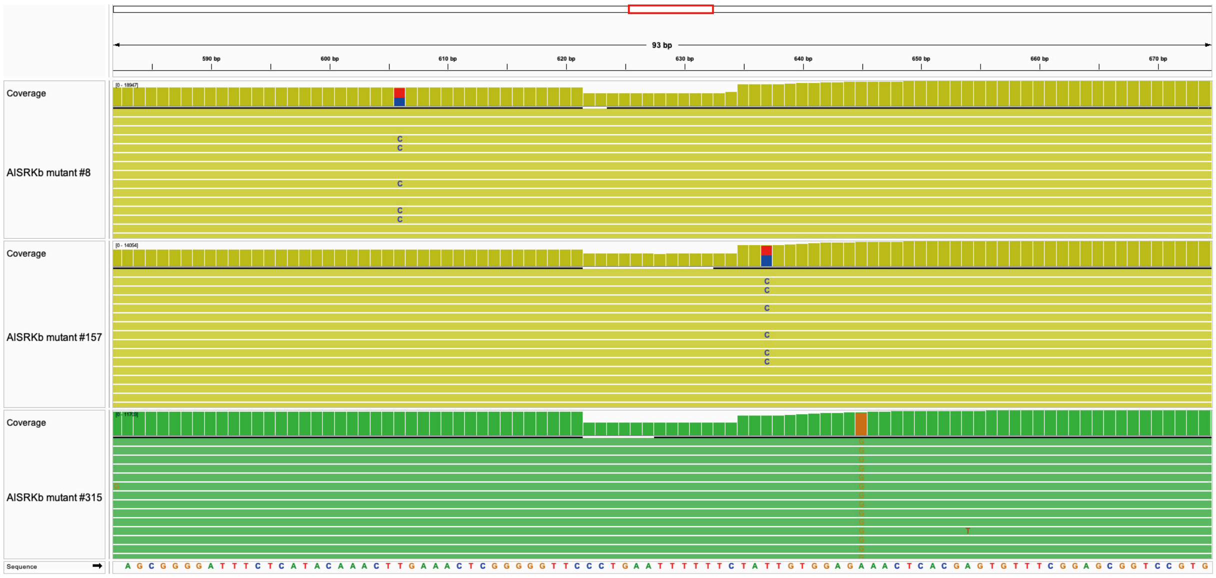

**Supplemental Figure S1.** Examples of mutated *AlSRKb* DNA sequences.
The DNA sequences of mutated *AlSRKb* genes from *A. thaliana* transformants #8, 157, and 315 were determined by amplicon sequencing and the nucleotide region between base pairs 582 and 674 of *AlSRKb* were visualized using the Integrative Genomics Viewer (IGV). The mutation A645G was observed in almost all *AlSRKb* genes in transformant #315 (shown in green). By contrast, the mutations T606C and T637C were observed in approximately 50% of *AlSRKb* genes in transformants #8 and #157 (shown in yellow).

Fig S2 Yamamoto et al.

**A**

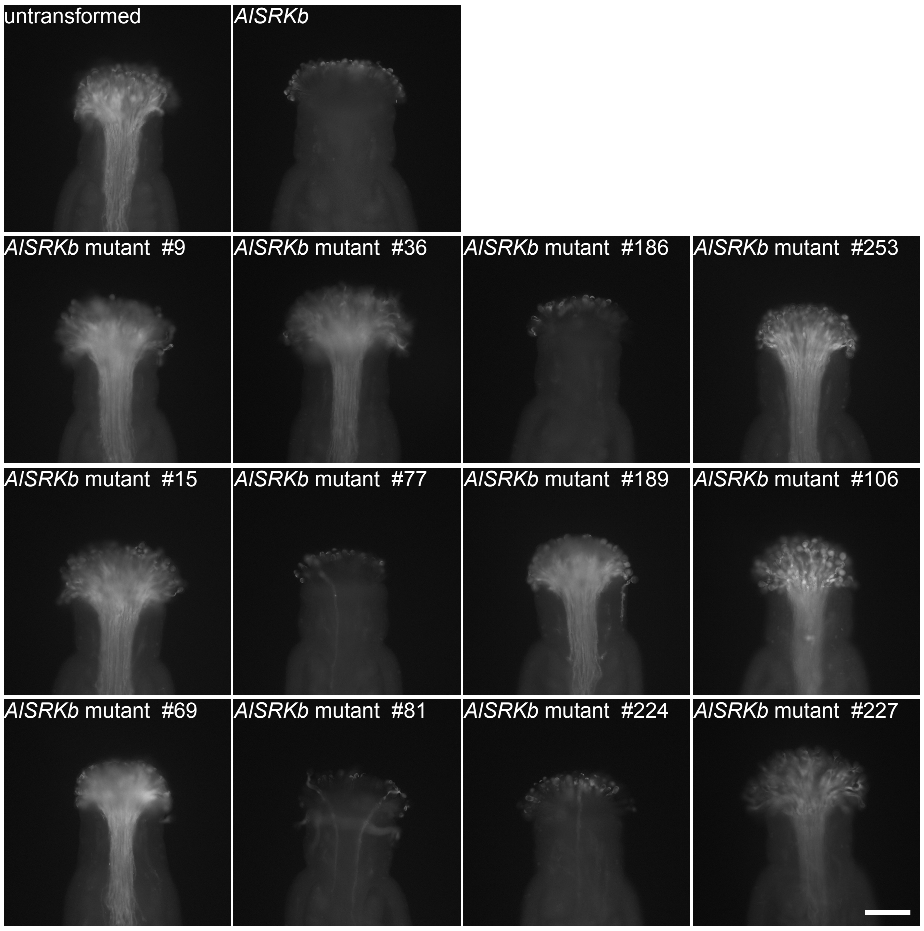


**Supplemental Figure S2.** Phenotypic analysis of *A. thaliana* transformants expressing mutated *AlSRKb-FLAG*.
**(A)** Microscopic observations were performed as described in Figure 1. Note that no or few pollen tubes germinated from AlSCRb-expressing pollen on the stigmas of transformants expressing mutated *AlSRKb-FLAG* genes #186, #77, #81, and #224, indicating that these transformants showed an SI response. Scale bar = 100 μm.
**(B)** Level of protein expression and the percentage of the endoglycosidase H-resistant form of mutated AlSRKb proteins in freshly constructed individual *A. thaliana* transformants expressing specific forms of mutated *AlSRKb-FLAG* genes. Experiments were performed and analyzed as shown in Figure 2A.

Fig S2 Yamamoto et al.

**B**

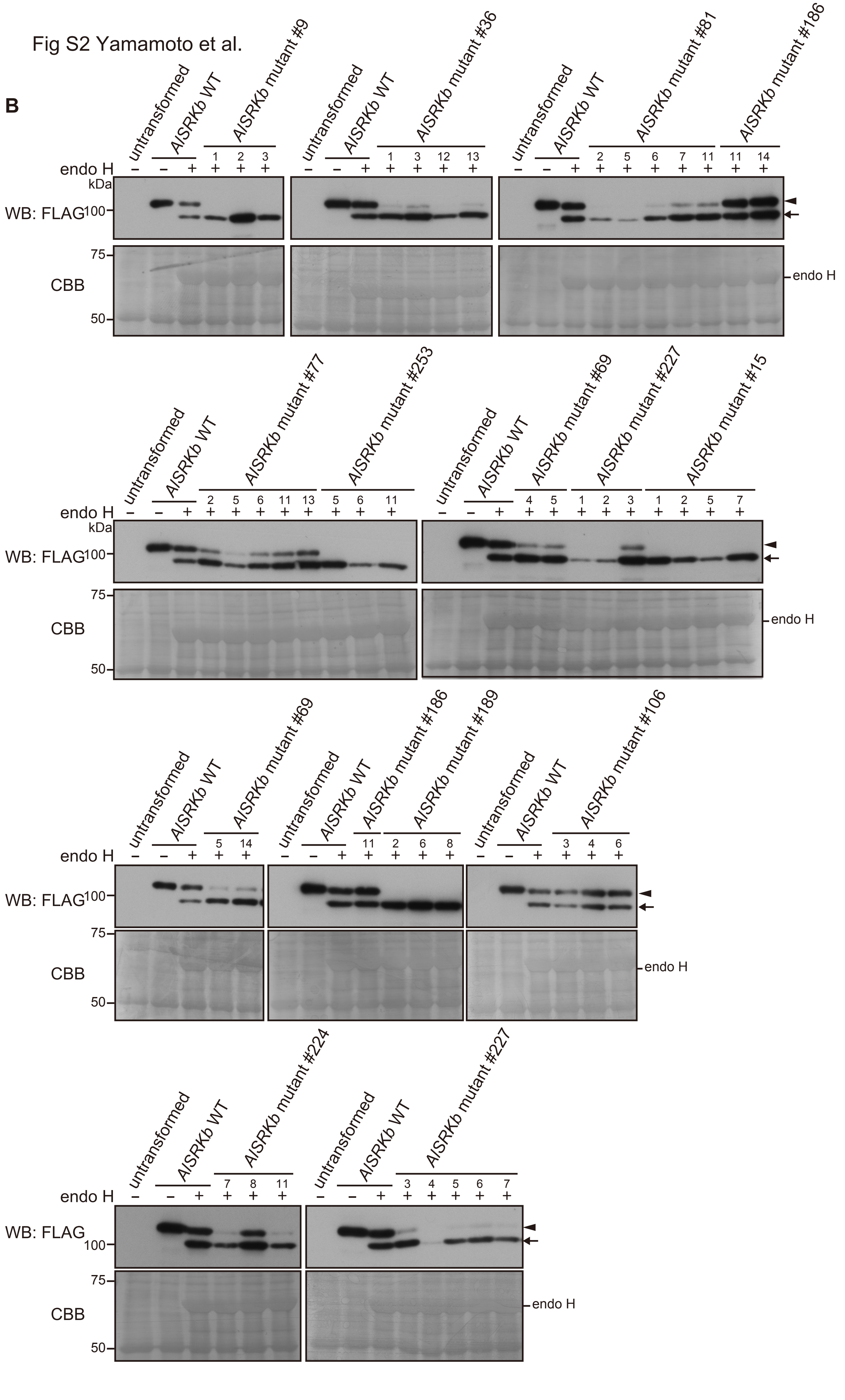

Fig S3 Yamamoto et al.

```
AlSRKb(S-20)   SYVHTLSSTESLTISSKQTIVSPGEVFELGFFNPAATSRDGDRWYLGIWFKTNLERTYVW
AlSRK01        IHANTLSSTESLTISRNLTIVSPGKIFELGFFKPST----RPRWYLGIWYKKIPERTYVW
AlSRK04        ------------------------------------------------------------
AlSRK05        ISANTLSVTESLTISSNQTIVSPGGVFELGFFKIL-----GDSWYLGIWYKKIPQRTYVW
AlSRK06        IYVNTLSPTESLTIASNRTIVSLGDDFELGFFKPAASLREGDRWYLGIWYKTIPVRTYVW
AlSRK09        ISVNTLSSTETLTISSNRTIVSPGYDFELGFFKPGS----SSLWYLGIWYKKVPDRIYPW
AlSRK13        ISTNTLSATESLTISSNKTIVSLGDVFELGFFTIL-----GDSWYLGIWYKKIPEKTYVW
AlSRK14        IYANTLLSTESLTIASNQTIVSLGDDFELGFFKPAASLRDGDRWYLGIWYKTISIRTYVW
AlSRK08        ------------------------------------------------------------
AlSRK15        ISVNTLSSTESLTISSNRTIVSPGGVFELGFFKPVS----NSRWYLGIWYKKVPQRAYVW
AlSRK16        ISANTLSATESLTISSNKTIVSPGGVFELGFFKIL-----GDSWYLGIWYKNVSEKTYVW
AlSRK18        INVNTLSSTESLTISSNRTIVSLGDDFELGFFKPAASLRNGDHWYLGIWYKTISVRTYVW
AlSRK19        ------------------------------------------------------------
AlSRK22        IYVNTLPSTEILTISSNRTIVSPGDVFELGFFKLGS----PARWYLGIWYKKVPEISYVW
AlSRK23        ------------------------------------------------------------
AlSRK25        ISANTLSATESLTISSNKTIVSPGGVFELGFFKIL-----GDSWYLGIWYKNVSEKTYVW
AlSRK30        ------------------------------------------------------------
AlSRK31        INVNTFSSSNSLTLSNNRTIVSSSDVFELGFFKITTSSPDDDRWYLGIWYKKIPERTYVW
AlSRK36        ISGNTLSSTETLTISSNRTIVSPGNDFELGFFKFDS----RSLWYLGIWYKKVPQRTYPW
AlSRK37        ISANTLSATESLTISSNKTIVSPGGVFELGFFRIL-----GDSWYLGIWYKKISQRTYVW
AlSRK38        INVNILSSTESLTISNNRTIVSPGGLFELGFFKPGT----SSRWYLGIWYKKIPEEAFVW
AlSRK39        INVNTFSSSNSLTLFNNRTIVSSSDVFELGFFKITSSSPDDDRWYLGIWYKKIPERTYVW
AlSRK50        ISVNTLSSTETLTISSNRTIVSPGDDFELGFFKTGS----SSLWYLGIWYKKVPDRTYVW
AlSRK52        ------------------------------------------------------------
AhSRK03        IFVNTLSSTESLTIASNRTIVSLGDDFELGFFRPAASLREGDRWYLGIWYKTISVRTYVW
AhSRK12        ISVNTLSSTESLTISSNRTIVSPSGVFELGFFETAP----NSRWYLGIWYKKVPEKTYVW
AhSRK13        INVNTLSSTESLTISSNRTIVSLGDVFELGFFNPTPSSRDGDRWYLGIWYKEIPKRTYVW
AhSRK15        ININILSSTESLTISSNRTIVSPGDVFELGFFKPAT----SSRWYLGIWYKTMPEGTYVW
AhSRK20        ISVNTLSSTETLTISSNRTIVSPGDDFELGFFKPGS----SSLWYLGIWYKKVPDRTYVW
AhSRK28        IFANTLSSTESLTIASNQTIVSLGDDFELGFFRPAASLREGDRWYLGIWYKTISVRTYVW
AhSRK32        ISVNTLSSTETLTISSNRTIVSPGDHFELGFFKTGS----SSLWYLGIWYKKVPQRTYAW
AhSRK43        STVNTLSATESLTISSNRTIVSPNDVFELGFFKPGT----SSRWYLGIWYKTILQRTYVW
AhSRK47        ISVNTLSSTESLTISSNRTIVSPGGVFELGFFETVS----TSRWYLGIWYKKVPQRTYVW
CgSRK1SD       -------------------------------------------------------RTYAW
CgSRK2SD       -------------------------------------------------------RTYAW
CgSRK3SD       -------------------------------------------------------RTYAW
CgSRK4SD       -------------------------------------------------------RTYAW
CgSRK5SD       -------------------------------------------------------RTYAW
CgSRK6SD       -------------------------------------------------------RTYAW
CgSRK7         ISANTLSATDSLT--SNKTLVSPGDVFELGFFKIL-----SDSWYLGIWYKTLPQKTYVW
CgSRK9SD       ------------------------------------------------------------
CgSRK11SD      ------------------------------------------------------------
CgSRK18SD      ------------------------------------------------------------
Cg2-2-KS2      TSVNTLSSTESLTISNNRTIVSPNGVFELGFFETAS----NSRWYLGIWYKKVPERTYVW
CgSRK34SD      ------------------------------------------------------------
CgSRK35        -----METTFCLSVS-KQTIVSPDDDFELGFFKTRS----SSLWYLGIWYKKVPDRTYVW
CgSRK36SD      -------------------------------------------------------RTYAW
CgSRK37SD      ------------------------------------------------------------
CgSRK47SD      -------------------------------------------------------RTYAW
```

**Supplemental Figure S3.** Alignment of amino acid sequences of the SRK variants from *Arabidopsis lyrata* (AlSRKx), *A. halleri* (AhSRKx), and *Capsella grandiflora* (CgSRKx) used in this study.

```
AlSRKb(S-20)  VANRDNPLYNSTGTLKISDTNLVLLDQFDTLVWSTNL-T-GVLRSP-VVAELLSNGNLVL
AlSRK01       VANRDTPLSNSVGTLKISDGNLVILDHSNIPIWSTNT-K-GDVRSP-IVAELLDTGNLVI
AlSRK04       -------------------GNLVILGHSDKLVWSTNL-TSGNVRSL-VVAELLANGNFVM
AlSRK05       IANRDNPLSNSIGILKISNANLVILDNSDTPVWSTNL-T-GAVRSL-VVAELLDNGNFVL
AlSRK06       VANRDNPLSSSAGTLKISGINLVLLNQSNITVWSTNL-T-GAVRSQ-VVAELLPNGNFVL
AlSRK09       VANRDNPLSNSLGTLRVSGTNLVLLDHSDKPVWSTNL-TTGNVKSP-VVAELLANGNFVL
AlSRK13       VANRDNPISTSTGILKISNANLVLLNHFDTPVWSTNL-T-AEVKSP-VVAELLDNGNFVL
AlSRK14       VANRDHPLYSSAGTLKISGINLVLLNQSNIAVWSTNL-T-GAVRSP-PVAELLPNGNFVL
AlSRK08       -------------------------NQSNITVWSTNL-T-GAVRPP-IVAELLPNGNFVL
AlSRK15       VANRDSPLPNSIGILKISDANLVLLDHSNTLVWSTNR-T-RDTRSP-MVAELLDNGNFVL
AlSRK16       VANRDNPLSDSIGILKITNSNLVLINHSDTPIWSTNL-T-GAVISP-VVAELLDNGNFVL
AlSRK18       VANRNHPISSSAGTLKISGINLVLLNQSNITVWSTNL-T-GAVRSP-VVAELLSNGNFVL
AlSRK19       ------------------------------------------------------------
AlSRK22       VANRDNPLSNSMGGLKIVDGNLIIFDHYDNYVWSTKL-TTKDVRSS-LVAELLDNGNFVL
AlSRK23       ---------------------------------------------------LLANGNFVL
AlSRK25       VANRDKPLSNSIGILKITNANLVLLNHYDTPVWSTNL-T-GAVRSP-VVAELHDNGNFVL
AlSRK30       ------------------------------------------------------------
AlSRK31       VANRDDPLSTSSGTLKISDNKLLLLDQVDTPIVSWNL-SGGGVRSP-VVAELLGNGNFVV
AlSRK36       VANRDNPLSNPIGTLKISGNNLVLLDHSNKPVWSTNL-TIRNVRSP-VVAELLANGNFVM
AlSRK37       VANRDNPLSNPIGILKISNANLVILDNSDISVWTTNL-T-GAVRSP-VVAELLDNGNFVL
AlSRK38       VANRDSPLFNAIGTLKISDTNLVLLDHSSTPVWSTNLSTRGVVRSS-VVAELLANGNFVL
AlSRK39       VANRDDPLSTSSGTLKISDNKLLLLDQVDTPIVSWNL-SGGGVRSP-VVAELLDNGNFVV
AlSRK50       VANRDNPLSEPIGTLKISGNNLVLLDHSNKLVWSTNL-TRGSMRSP-VVAELLANGNFVM
AlSRK52       ------------------------------------------------------------
AhSRK03       VANRDHPISSSDGTFKISGINLVLLNQSNMTVWSTNL-T-GAVRSP-VVAELLANGNFVL
AhSRK12       VANRDHPFSNSIGILKISEANLVLLDHSDTLVWSTNR-T-GGTRSP-VVAELLDNGNFVL
AhSRK13       VANRDNPLSNSTGTLKISDNNLVLVDQFNTLVWSTNV-T-GAVRSL-VVAELLANGNLVL
AhSRK15       VANRGNPLSSSIGTLKIVGSNLVLLADSDIPIWSTNL-TGGDVRST-VVAELLANGNLVL
AhSRK20       VANRDNPLSNSTGTLKISGNNLVLFGRSDKPVWSTNL-TRGNVRPP-VMAELLANGNFVI
AhSRK28       VANRDHPISSSDGTLKISGINLVLLNQSNITVWSTNL-T-GAVRSP-VVAELLPNGNFVL
AhSRK32       VANRNNPLSNSIGTLKISGSNLVLFGHSNKPVWSTNL-TRGNVRSP-VMAELLANGNFVM
AhSRK43       VANRDKPLINPIGTLKISNTNLVLLDSSDTLVWSTNL-TERDVISP-VVAQLLDNGNFVL
AhSRK47       VANRDNPLSNSIGILKILDANLVLLDHSDTLVWSTNR-T-GDTKSP-LLGELFDNGNFVL
CgSRK1SD      VANRDHPISSSDGTLKISGINLVLLNQSNITVWSTNL-T-RAVRSP-VVAELLGNGNFVL
CgSRK2SD      VANRDNPLSSSIGTLKILDSNLMLLDQSDTTVWSTNL-T-GAVSSS-VVAELLSNGNFVL
CgSRK3SD      VANRDTPLSDSVGTLKISKGNLVIFDHSNIPIWSTKT-I-GDVRSP-IVAELLDTGNLVL
CgSRK4SD      VANRDNPLSNSAGTLKISGINLVLLNQSNISVWSTNL-T-GAMRSP-VVAELLSNGNFVL
CgSRK5SD      VANRDNPLSSFAGSLRFSGINLVLLNQSNISVWSTNV-T-GAVRSA-VVAELLSNGNFVL
CgSRK6SD      VANRDKPLYNSAGTLKISGINLVLLDQSNITVWSTNL-T-GAETSP-VVAELLPNGNFVL
CgSRK7        IANRDNPLFGSTGVLKISNANLILQSQTDTLVWSTNL-T-GAVRAP-MVAELLDNGNFVL
CgSRK9SD      ------------------------------------------------------------
CgSRK11SD     ------------------------------------------------------------
CgSRK18SD     ------------------------------------------------------------
Cg2-2-KS2     VANRDNPFSTSIGTLKISEANLVLHDHSNTIVWSTNR-T-GDTRSP-VVAELLDTGNFVV
CgSRK34SD     ---------------------------------------------P-LVAELLPNGNFVL
CgSRK35       VANRDNPLSNPLGTLKISGNNLALLDHSNKPVWSTNL-TRGNMRSP-VVAELLANGNFVL
CgSRK36SD     VANRDNPLPSSIGTLKISNMNLVLLDHSNKSVWSTNL-TRGNERSA-VVAELLANGNFVM
CgSRK37SD     ------------------------------------------------------------
CgSRK47SD     VANRDNPLPSSTGTLKISNMNLVLLDYCNKSVWSTNL-TRGNERSSPVVAELLANGNFLM
```

Accession numbers: AlSRK01(KJ772402.1), AlSRK04,(AAK19315.2), AlSRK05(AIB04258.1), AlSRK06(ACU29642.1), AlSRK08(AAO16816.1), AlSRK09(ABF71372.1), AlSRK13(BAB40986.1), AlSRK14(AID21648.1), AlSRK15(AGC54823.1), AlSRK16(ADQ37362.1), AlSRK18(AID21658.1), AlSRK19(AAK19319.2), AlSRK22(ABF71375.1), AlSRK23(AAK19318.2), AlSRK25(ACU29643.1), AlSRK30(ACP31395.1), AlSRK31(ABF71377.1), AlSRK36(AGC55017.1), AlSRK37(ABF71379.1), AlSRK38(ADQ37372.1), AlSRK39(AID21691.1), AlSRK50(ADQ37382.1), AlSRK52(APC45346.1), AhSRK03(AID21586.1), AhSRK12(AID21577.1), AhSRK13(AJP61130.1), AhSRK15(KJ772390.1), AhSRK20(AID21613.1), AhSRK28(AJP61126.1), AhSRK32(AJP61104.1), AhSRK43(AJP61150.1), AhSRK47(AFP33764.1), CgSRK1SD(ABF72159.1), CgSRK2SD(ABF72160.1), CgSRK3SD(ABF72161.1), CgSRK4SD(ABF72162.1), CgSRK5SD(ABF72163.1), CgSRK6SD(ABF72164.1), CgSRK7(ABP02072.1), CgSRK9SD(ACO83282.1), CgSRK11SD(ACO83283.1), CgSRK18SD(ACO83280.1), Cg2-2-KS2(VUE13615.1), CgSRK34SD(ACM47717.1), CgSRK35(SSC84616.1), CgSRK36SD(ACM47719.1), CgSRK37SD(ACO83286.1), CgSRK47SD(ACM47720.1)

```
AlSRKb(S-20)   KDSKTN-DKDGILWQSFDYPTDTLLPQMKMGWDVKKGLNRFLRSWKSQYDPSSGDFSYKL
AlSRK01        RYFNNN--SQEFLWQSFDFPTDTLLPEMKLGWDRKTGLNRFLRSYKSSNDPTSGSFSYKL
AlSRK04        RYS-SK-DQGGFLWQSFDYPTDTLPPQMKLGWDRKTKLNRYLRSWKSSYDPSSGNFTYEL
AlSRK05        RNSKTN-DLDGFLWQSFDFPTDTLLPQMKLGLDNKRGLNKFITSWKSSFDPSSGDYIFKL
AlSRK06        RDSKSN-GQDVFFWQSFDHPTDTLLPHMKLGLDRKTENNRVLTSWKNSYDPSSGYLSYKL
AlSRK09        RYTNNN-DPSGFLWQSFDFPTDTLLPEMKLGYDLKTGVNRFLRSWRSFDDPSSGNFTYKL
AlSRK13        RDSKTN-GSDEFLWQSFDFPTDTLLPQMKLGLDHKKRLNKFLRSWKSSFDMSSGDYLFKI
AlSRK14        RYSKTN-GQDILLWQSFDYPTDTLLPHMKLGLDLKTGNNRLLTSWKNSFDPSSGYISYKL
AlSRK08        RDSKTN-GQDGFLWQSFDYPTDTLLPHMKLGLDRKTGNNRFLTSWKNSYDPSSGSLSYKL
AlSRK15        RVSSNKNDQGGILWQSFDFPTDTLFPQMKLGWDLKAGINRYLISWKSPNDPSSGKYSYKL
AlSRK16        RDSKTN-DSDGFLWQSFDFPTNTLLPQMKLGLDNKRALNRFLTSWKNSFDPSSGDYTFKL
AlSRK18        RDSKPN-EQDRLLWQSFDYPTDTLLPHMKLGLDLKTGNNRFITSWKNSYDPSSGYLSNKL
AlSRK19        --------------------------------------------WKNSYAPSSGDLSYKL
AlSRK22        RVSNNN-DPDKFLWQSFDYPTDTLLPQMKLGWDLKTGLNRFLRSWKSSDDPSSGNFTCKL
AlSRK23        RESGNK-DQDGLVWQSFDFPTDTLLPQMKLGWDRKTGLNKILRSWKSPSDPSSGYYSYKL
AlSRK25        RDSKTN-ASDRFLWQSFDFPTNTLLPQMKLGWDHKRGLNRFLTCWKNSFDPSSGDYMFRL
AlSRK30        ------------------------------------------------------------
AlSRK31        KDSKAN-NPNGFLWQSFDFPTDTLLPQMKMGWDRKTANNRFLRSWKSPYDPSSGDYSYKL
AlSRK36        RYS-NN-DQGGFLWQSFDYPTDTLLPQMKLGWDRKTGLNRILRSWRSLDDPSSSNYSYEL
AlSRK37        RDSKIN-ESDEFLWQSFDFPTDTLLPQMKLGQDHKRGLNRFLTSWKSSFDPSSGSFMFKL
AlSRK38        RYSNNS-DPSGFLWQSFHFPTDTLLPQMKLGWDRKTGRNTFLRSWRSPDDPSSGAFSYKL
AlSRK39        KESKAN-NPNGFLWQSFDFPTDTLLPQMKMGWDRKTANNRFLRSWKSPYDPSSGDYSYKL
AlSRK50        RYY-NN-DRGVFLWQSFDYPTDTLLPQMKLGWDRKTGLNRFLRSSKSLDDPSSGNFSYKL
AlSRK52        ----------------------TLLPQMKLGRDFKTGLNKFLTSWKSQIDPSSGNFSYKL
AhSRK03        RDSKTN-GKNGFLWQSFDYPTDTLLPHMKLGLNVKTKNNRFLTSWKNSYDPSSGSFSYKL
AhSRK12        RESSNKNDLDRYLWQSFDFPTDTLLPEMKLGWDLKTGLNRYLTSWKSPNDPSSGYYSYKL
AhSRK13        RDSKIN-ETDGFLWQSFDFPTDTLLPEMKLGWDLKTGVNKFLRSWKSPYDPSSGDFSYKL
AhSRK15        RHSNKN-KSGEFLWQSFDFPTDTLLPEMKLGWDRKTGFNRFLRSWKSPDDPSEGNFLYKL
AhSRK20        RYS-KN-DQGGFLWQSFDFPTDTLLPQMKLGWDLKRGLNRFLRSWKSLDDPSSGDFSYKL
AhSRK28        RNSKTN-GHDVFMWQSFDNPTDTLLPHMKLGLDLKTGNNRFLTSWKNAYDPSSGYLSYKL
AhSRK32        RYSNNN--QGGFLWQSFDYPTDTLLPQMKLGWDHKTGLNRFLRSWKSVDDPSSGEFSYKL
AhSRK43        RYSNKD-VQSEFLWQSFHFPTDTLLPQMKIGLDRKTEFNRFLRSWRSADDPASGDYSFKL
AhSRK47        RESNNKNDQDGLLWQSFDFPTDTLLPQMKLGWDRKTGRNKFLISWKSPSDPSSGYYSYKL
CgSRK1SD       RDSKSN-GKDRFFWQSFDYPTDTLLPHMKLGFDLKTKNNRFLTSWRNAYDPSSGSFAYEL
CgSRK2SD       RDAKTN-DPDVFLWQSFDFPTDTLLPHMKLGWDLKTGRHRSLKSWRSLYDPSSGDLSYKL
CgSRK3SD       RYSNKN--SREFLWQSFDFPTDTLLPEMKLGWDRKTGLNRLLRSYESSNDPTSGSFSYKL
CgSRK4SD       KDSKTN-GKGGLLWQSFDYPTDTLLPHMKLGLDLKTKNNRFLTSWKNSYDPSSGSLLYKL
CgSRK5SD       RDSNTN-GKDGLLWQSFDYPTDTLLPHMKLGLDLKTGHNRVLTSWKNSYDPSRGFYLFQL
CgSRK6SD       RGSKGN-GKDGFFWQSFDYPTDTLLPHMKLGLDLKTENNRFLTSWKNAYDPSSGSFSYKL
CgSRK7         RDSKTN-GSDGFLWQSFDFPTDTLLPQMKLGRDHKRKLDRFLTSWKSSFDLSNGDYLFKL
CgSRK9SD       -------------WQSFDYPTDTLLPDMKLGLDFKTGNNRFITSWKNSYDPSSGNLSYKL
CgSRK11SD      ---------------SFDFPTNTLLPQMKLGWDLKRGLNRFLTSWKNSFDPSSGVYMFKL
CgSRK18SD      -----------------DFPTDTLLPQMKLGRDFIRGLNKSLTSWKTSFDPSSGDYVFKV
Cg2-2-KS2      RESSNKNDLDRFLWQSFDFPTDTLLPQMKLGWDLKKRLNRYLISWKSPNDPSSGYYSYKL
CgSRK34SD      RDPNTN-RQNGFFWQSFDYPTDTLLPHMKLGLDRKTGNNRNLTSWKNSFDPSSGYLCYKL
CgSRK35        RYTNNN-DPSGFLWQSFDFPTDTLLPEMKLGCNLKTGHNKFLRSWRSRDDPSSGNYLYKL
CgSRK36SD      RDSNNK-DASGFLWQSFDYPTDTLLPEMKLGYDLKTGLNRFLTSWRSSDDPSSGDFLYEL
CgSRK37SD      -------------WQSFHFPTDTLLPQMKLGWDRKTGRNIFLRSWRSSDDPSTGKFSYRL
CgSRK47SD      R------DRSGVLWQSFDYPTDTLLPEMKLGYDLKTGRNRFLTSWRSSDDPSSGEFSYKL
```

```
AlSRKb(S-20)  ET--RGFPEFFLL-WRNS------RVFRSGPWDGLRFSGIPEMQQWEY--MVSNFTENRE
AlSRK01       ET--GVYSEFFML-AKNS------PVYRTGPWNGIQFIGMPEMRKSDY--VIYNFTENNE
AlSRK04       ET--RGFPEFFLR-KTDI------PIHRSGPWDGIRISGIPEERQVDY--MVYNFTEDRE
AlSRK05       ET--QGLPEFFIS-RSKF------KLFRSGPWDGNRFSGIPEMEQWDN--IIYNFTDNRE
AlSRK06       EM--LGLPEFFMW-RSKV------PVFRSGPWDGIRFSGIPEMQIWKHINISYNFTENTE
AlSRK09       DT--QGLPEFWFR-ESDF------RLQRSGPWDGIQFSGIPEVRQLNY--MSYNFTENRE
AlSRK13       ET--LGLPEFFIW-MSDF------RVFRSGPWNGIRFSGMLEMQKWDD--IIYNLTENKE
AlSRK14       ET--LGLPEFFMW-RNEV------PIFRSGPWDGTRLSGIPEMQRWKDINISYNFTENKE
AlSRK08       EI--QGLPEFFVS-KSGV------PVFRSGPWDGIQFSGIPEMQRWKHVNISYNFTENKE
AlSRK15       ES--QGLPEFFLY-NRDS------PTHRSGPWDGIRFSGIPDKQQLDY--MVYNFTENKE
AlSRK16       ET--RGLTELFGL-FTIL------ELYRSGPWDGRRFSGIPEMEQWDD--FIYNFTENRE
AlSRK18       DI--LGLPEFLVL-REGV------TVYRSGPWDGIQFSGIPEMQRWKDFNIVYNFTENKE
AlSRK19       EN--RVLPEFFIR-KRDS------PMFRSGPWDGIKFSGMPDMQQWSYINIVYDFTKNRE
AlSRK22       ET--RGFPEFLIR-FRFT------PIYRSGPWDGIRFSGMPEMRDLGY--MFNKFTANGE
AlSRK23       EF--QGLPEYFLN-NRDS------PTHRSGPWDGIRFSGIPE-KPLKY--MVYNFTENKE
AlSRK25       DT--QGLPEFFGL-KNFL------EVYRTGPWDGHRFSGIPEMQQWDD--IVYNFTENSE
AlSRK30       -----VLPEFFIW-QTDI------PTYRSGPWDGVRFSGMVQMRDLDY--MVNNFTDNRE
AlSRK31       EI--QGLPQFYLW-TAKR------AVFRSGPWDGIRFSGMPEMQRWNNAEIVYNFTDNRE
AlSRK36       QT--RGFPEFFLL-DEDV------PVHRSGPWDGIQFSGIPEVRQLNY--IINNFKENRD
AlSRK37       ET--LGLPEFFGF-TTFL------EVYRSGPWDGLRFSGIPEMQQWDD--IIYNFTENRD
AlSRK38       ET--RSFPEFFIW-NTDA------PMYRSGPWDGVRFNGMVEMKELGY--MVSNFTDNRE
AlSRK39       EI--QGLPQFYLW-TSNA------QVFRSGPWDGIRFSGMPEMQRWNNADIVYNFTDNRE
AlSRK50       ET--RGLPEFFLL-MNDV-----LKIHRSGPWDGTQISGIPEERKLDY--MVYNFTENRG
AlSRK52       EI--QGLPEFYLS-TRNG------PYHRSGPWDGIQFSGIPDNQQLNY--LVYSFTENKE
AhSRK03       EIPRHGLPEFLMW-RSGG------PAFRSGPWDGIRFSGIPEMERWKFVNIVYNFTENKE
AhSRK12       EL--QGLPEFFLS-YKDS------PMHRSGPWDGVRFSGMPEKKQLTY--MVYNFTENEE
AhSRK13       ET--REFPEFFLS-WSNS------PVYRSGPWEGFRFSGMPEMQQWTN--IISNFTENRE
AhSRK15       ET--RGSSEFYIL-KEGE------QMYRSGPWDGIRFNGMPEMQKLSF--MGYNFTENQE
AhSRK20       ET--RGFPEFYLW-KNDI------PIHRSGPWDGTRISGVPEEQQLNY--MVYNFTENRE
AhSRK28       EM--QGLPEFLML-RGGG------PVFRSGPWDGFRFSGIPEMQNWKFAYIVYNFTENKE
AhSRK32       ET--RGFPEFFLR-KNDI------PVHRSGPWDGIRISGIPEDGQLDY--MVYNFTENSE
AhSRK43       KT--RGVPEFFIWVKQNT------RMYRSGPWNGIRFSGMPEMLEFDY--MVYNFTENRE
AhSRK47       DF--QGIPEFFLN-NRGW------PTHRSGPWDGIRFSGIPE-KQLNY--MVYNFTENKE
CgSRK1SD      QIPKNGLPEFFML-RSGG------PALRSGSWDGFRLSGIPEMQRWSFLNIVYNFTENKE
CgSRK2SD      ET--RGLPDFFIW-KTDV------RVYRSGPWDGIRFSGIPEMPRWNF--IVNNFTENRE
CgSRK3SD      EI--GAYSEFFML-ADNS------PVYRSGPWNGIQFIGMPEMRKSDY--VVYNFTESDE
CgSRK4SD      EM--LGLPEFFMW-RSGG------LVFRSGPWDGFRFGGIPEMERWKFVNIVYNFTENKE
CgSRK5SD      QI--PGLPEFFLW-KSDF------LWFRSGPWDGIRFSGIPDMQQWLNFNFVYNFTENKE
CgSRK6SD      EVPISGLPEFFLW-RNGG------PVIRSGAWDGFRFTGIPEMQNWKLINIVSNFTDNKE
CgSRK7        ET--QGLPEFFLW-KKFW------ILYRSGPWDGSRFSGMSEIQQWDD--IIYNLTDNSE
CgSRK9SD      NI--LGLPEFVML-RDVV------TVFRSGPWDGIHFSGIPEMQTWKDINIAYNFTENKE
CgSRK11SD     ET--QGLPDFFGL-YGIW------KMYRSGPWDGFRFSGIPEMQQWND--IIYNFTENKE
CgSRK18SD     EP--QGIPEFFTWKKRMF------RLFRTGPWDGIGFSGIPDMHLWDD--LIYNFTENTV
Cg2-2-KS2     EL--QGLPEFFLS-YKDS------PMHRSGPWDGIRFNGIPEKQQLTY--MVYNFTENEE
CgSRK34SD     EI--LGLPEFLMW-KDES------PMIRSGPWDGTRFSGIPDMQSWKIANVVYNFTDNKE
CgSRK35       KT--QGLPECFLL-DDSVKGAKAVKVHRSGPWNGIQFSGLPEVQQLEY--MDYKFTENRD
CgSRK36SD     EA--RRLPEFYLS-NGIF------RLYRSGPWNGIRLSGIRDDQKLSY--LVYNFTENSE
CgSRK37SD     ET--RSFPEFFIW-QTDV------PMYRSGPWDGVRFSGMVEMRDLDY--MVYNFTDNQE
CgSRK47SD     DN-KMGLPEFYLF-KDDF------RVHRSGPWNGIRFSGIPDDQSLSY--MVYNFTENSQ
```

```
AlSRKb(S-20)   EVAYTFQITNHNIYSRFTMSSTGALKRFRWISSSEEWNQLWNKPND-HCDMYKRCGP-YS
AlSRK01        EVSLTFLMTSQNTYSRLKLSDKGEFERFTWIPTSSQWSLSWSSPKD-QCDVYDLCGP-YS
AlSRK04        EVAYTFLVTNHSIYSRLTMSYSGYFQRFTWTSPAWGWRQLWSAPMDLQCDLYPSCGP-YA
AlSRK05        EVAYTFQLTNHSLYSRLKINSDGLLQRFTWIPTIQEWNMYWLTPAT-HCDFYENCGP-YA
AlSRK06        EVAYTYRVTTPNVYARLMMDFQGFLQLSTWNPAMSEWNMFWLSSTD-ECDTYPSCNPTNS
AlSRK09        EVTDTFLMTNHSIYSRLTVSAAGSFDRFTWITPSTGWSRYWSLPTD-ECDSFKSCGP-YA
AlSRK13        EVAFTFRPTDHNLYSRLTINYAGLLQQFTWDPIYKEWNMLWSTSTDNACETYNPCGP-YA
AlSRK14        EVAFTFRVTTPNVYSRLIMNSEGFLQLSRWNPTLSEWNVFWRSSTS-DCNGYQSCTP-YS
AlSRK08        EVAFTYRVTIPDAYAGMTMDSEGLLQLFTWIPTTLEWNMFWLSSAG-ECDIYQRCSP-YT
AlSRK15        EVAYMFSMTNHSIYSRLTLSSIGTFERFTWIPPSWQWNLLWSSPKH-ECDAYERCGP-YS
AlSRK16        EVFYTFRLTDPNLYSRLTINSAGNLERFTWDPTREEWNRFWFMPKD-DCDMHGICGP-YA
AlSRK18        EIAFTYRVTTPKVYARLTMNFDGYLQLSRWLPETLEWNVFWQTSAA-DCEVYMSCTP-NS
AlSRK19        EATYSYRPTNQNIYSRLTMSSSGLLQLFTWSSTSQEWNLVWFSSKD-ECDIYGICSP-NA
AlSRK22        EVAYTFLMTNKSIYSRITLSSAGIFERYTWVPTSWEWTLFSSSPTD-QCDMNEECGP-YS
AlSRK23        EVAYTFTMIDHSIYSRLTVSPTGTLNRFTMIPPSWQWNMVWFSPKD-ECDMYETCGP-YG
AlSRK25        EVAYTFRLTDQTLYSRFTINSVGQLERFTWSPTQQEWNMFWSMPHE-ECDVYGTCGP-YA
AlSRK30        EVVYKFLMTDNHILSRLTLSPLGYLQQITW--KYEDRILSWLSPTD-PCDAYQICGP-YS
AlSRK31        ETAFTFQDIDPSSYSRLKMSFSGLLELSTWVPTTLAWDNFWLLSTN-PCDMFEVCGS-YS
AlSRK36        EISYTFQMTNHSIYSRLTVSFSGSLKRFMYIPPSYGWNQFWSIPTD-DCDMYLGCGP-YG
AlSRK37        EVAYTFRVTEHNSYSRLTINTVGRLEGFMWEPTQQEWNMFWFMPKD-TCDLYGICGP-YA
AlSRK38        EIAYTFQMTKHHIYSRLTMSPTGYLQQITFIEKNENRILSWFSPMD-QCDVYKVCGP-YS
AlSRK39        EIAFTFRDADPSSYSRLKMSTLGLLELSTWVPTTPGWKNFWISSIN-PCDMYEECGP-YS
AlSRK50        EVVYKFLMTNHSIYSRLILSNLGYLQRFTWFPPSWGWIQFWSSPRDFQCDLYQTCGP-YS
AlSRK52        EVTYTFSMTNHSIHSRLTVRTDGTLIRFTWIPKFTGWSMVWSAPKD-DCDFKASCGS-YS
AhSRK03        DIAFTFRVTTPDVYAKLTMRFDGFLELSTWDPEMLEWNVFWVTSTS-DCDIYMGCTP-YS
AhSRK12        EVAYTFSMTNHSILSRLTVSSSGTFNRFTWIPPSWQWNTVWFSPKD-DCDLYERCGP-YS
AhSRK13        EIAYTFRDTDQNIYSRLTMSSSGYLQRFKWISNGEDWNQHWYAPKD-RCDMYKKCGP-YG
AhSRK15        EVTYTFLMTNHSIYSRLTLTPSGSLQQFTWIPTERENDLFWNSPKD-QCDAYEKCGP-YS
AhSRK20        EVAYTFLMTNSSIYSRLTLSHSGFFQRLTSIPPAWGWSVLWSSPMDRPCDYYQFCGP-YS
AhSRK28        DVAFTYRVTTPNFYAKLTMRFEGFLELSTWDPDMLEWNVFWVSSTA-DCNIYMGCTA-NS
AhSRK32        EVSYTFLVTNHSLYSRLRMSSSGYFQRFTYTPPAWGWRQLWSAPMDLQCDLYQKCGP-YA
AhSRK43        EIVYTFLMTNHSIYSRLTMTPAGYLQQSTWFPTEEE--ASWVSPNE-QCDTYRICGP-YG
AhSRK47        EVTYTFSMINHSIYSRLTMNPTGTFSRFTWIPTSWQWSVPWFSPKD-ECDMYKTCGS-YG
CgSRK1SD       DVAFTYSITTPNVYAKLTMKFDGFLELSSWDPEMLEWNVFWVSSTT-DCDTYMGCTA-YS
CgSRK2SD       EITYSYRVTDHNTYSRLILSSSGVLQQFTWSPNEQEWSMFWTSPKD-LCDTYRKCGP-YS
CgSRK3SD       EVSFTFQMTNQNTYSRLTLNHEGEFARFTWIPTSSQWSLSWSSPKD-QCDVYDLCGP-YS
CgSRK4SD       EIAFTYRVTTPNVYARMMMNFDGFLTTMTWIPDTLEWNIVWQTSAD-SCDVYMSCTP-NS
CgSRK5SD       EVAYTYRVTTPNTYSRLTLNSEGILQLFTWLPETLEWNMVWMSYLA-ACDLYRVCSR-YS
CgSRK6SD       DVAFTYRVTTPNVYAKLTMKFDGFLELSTWDPEMSEWNVFWVSSSG-DCDTYMWCTT-NS
CgSRK7         EVAFTFRLTDHNLYSRLTINDAGLLQQFTWDSTNQEWNMLWSTPKE-KCDYYDPCGP-YA
CgSRK9SD       EIAFTYRVTVPNVYAKLTMNFDGFLQLSSWIPETLEWNVFWQTSQG-DCDVYMSCTP-NS
CgSRK11SD      EIAYTFRLTDTNFYSRLTINSVGSLERFTWDPTRQEWNRFWTMPKD-DCDTHGICGP-YA
CgSRK18SD      EVAYSFRLTNHSLYSRLTINSDGLLQRFEWIPEDQEWTIFWSTLKE-SCDIYNSCGP-YA
Cg2-2-KS2      EVAYTFSMTNQSILSRLTVSSSGTFDRFTWVPPSWQWNTVWSSPKD-ECDLYKTCGP-YS
CgSRK34SD      EVAFTYRVTTPNVYSRLTMNSDGFLQLFTWDRTMSEWSLFWLSSVS-ECDAYQICTL-FS
CgSRK35        EVAYTFLMTNHSIYSRVTISDSGALHRYTWIPPSYGWNLFWTTPTD-QCEMYKVCGP-YG
CgSRK36SD      EVAYAFQMTNNSIYSKITLSVSGNFERQTWNPSLGMWNVFWSFPLDSQCDTYRICGP-YS
CgSRK37SD      EVVYTFLMTNHDIYSRLTMSPAGSLQQITW--KDEDRILSWLSPTD-PCDAYQICGP-YS
CgSRK47SD      EVAYTFRMTNNSIYSRLTLSSEGYIERLTWNPSSGVWILYWSSPFHSQCDMYKMCGA-YA
```

```
AlSRKb(S-20)   YCDMNTSPICNCIGGFKPRNLHE-WTLRNGSIGCVRKTRLNC-GGDGFLCLRKMKLPD-S
AlSRK01        YCDINTSPICHCIQGFEPKFPE--WKLIDVAGGCVRRTPLNC-GKDRFLPLKQMKLPD-T
AlSRK04        YCDTNTPPLCNCIRGFNPWSMEQ-WNMGDGTSGCVRRTPLSC-RRDGFLPMKKMKLPT-T
AlSRK05        YCDMNTSPMCNCIQGFEPRIPQE-WASGDVAGSCQRKTPLNC-LEDGFIKLKKMKLPA-S
AlSRK06        YCDANKMPRCNCIKGFVPGNPQE-RSLNNSFTECLRKTQLSC-SGDGFFLMRKMKLPA-T
AlSRK09        YCDLNTSPVCNCIGGFDPKNQQE-WDLREGGTGCVRRTPLSCTGDDGFLKLKNMKLPD-T
AlSRK13        YCDMSTSPMCNCVEGFKPRNPQE-WALGDVRGRCQRTTPLNC-GRDGFTQLRKIKLPD-T
AlSRK14        YCDTNTTPNCNCIKGFAPQNPQE-GALDNTNTECVRKTQLSC-DGDGFFWLRNMKPPD-T
AlSRK08        YCDRNKTPNCNCIKGFEPMDPLE-EARDNTYIECIRKTQLSC-SGDRFFRLSKMKVPD-T
AlSRK15        YCDVNTSPICNCIEGFDPMNQQQ-WDLSNGAGGCLRRTQLSC-TGDGFLRLQKMKLPDTV
AlSRK16        YCDTSTSPACNCIRGFQPLSPQE-WASGDASGRCRRNRQLNC-GGDKFLQLMNMKLPD-T
AlSRK18        YCDPTKTTKCNCIKGFEPRDPRE-GALDTTNTDCVRKTQLSC-NGDGFFWLRNITPPD-T
AlSRK19        YCDVSTSPACNCIRGFQPRNEQN-WVLNGGSGECVRKTQLSCNGGDGFYQLKKVKLPD-T
AlSRK22        YCDTSTSPVCNCIQGFSPRSQQQ-WDLADGLSGCVRRTPLSC-RGDRFLRLKNMKLPD-T
AlSRK23        YCDINTSPTCNCIKGFDPKYQQQ-WDLSNGVGGCVRRTPLNC-SEDGFVQLKNMKLPD-T
AlSRK25        YCDMSKSPACNCIKGFQPLNQQE-WESGDESGRCRRKTRLNC-RGDGFFKLMNMKLPD-T
AlSRK30        YCYLKTSAFCNCIKGFEPKIPEA-WAVSDGTSGCVRKTRLSCSRGDVFFQLKNTKLPD-T
AlSRK31        YCDTNTSPMCNCFKGFDPMNPHD-WYSGDWSSGCVRKNPLSC-TGDGFLQLKKMKLPDTT
AlSRK36        YCDVNTSPICNCIRGFEPRNLQE-WILRDGSDGCVRKTQLSC-GGDGFVELKKIKLPD-T
AlSRK37        YCDMSTSPACNCIKGFQPLSQQE-WASGDVTGRCRRKTQLTC-GEDRFFKLMNMKLPA-T
AlSRK38        YCYMSTSPLCNCIQGFEPKIWRA-WELKDGTSGCVRKTRLSCGSGDGFLRLEKMKLPN-T
AlSRK39        YCDTNTLPMCNCIKGFDPMNSDE-WNSKDGSSGCVRRTPLSC-KEKEFVQLKKMKLPD-T
AlSRK50        YCDMNTLPLCNCIRGFRPWNEQQ-WELRDGSSGCVRKTPLSC-DGDGFWRLKNMKMPD-T
AlSRK52        YCDQNTSPICNCIKGFKPFYDQQVWELDGWTAGCVRRTKLSC-RGDGFWRLTKMKLPD-T
AhSRK03        FCDMNTTPKCNCIKGFEPSNPQG-GAMNNTSTECVRKTQLNC-KGDGFYWLRNMKLPE-T
AhSRK12        YCDVNTSPSCNCIQGFDPKNQQQ-WDLSNGVSGCVRRTRLSC-SEKRFLRLKKMKLPV-T
AhSRK13        ICDTNSSPECNCIKGFQPRNLQE-WSLRDGSKGCVRKTRLSC-SEDAFFWLKNMKLPD-T
AhSRK15        YCNMFTSSMCNCIKGFEPKNPQE--ALTDGLDGCVRKTKLSC-TGDGFWKLSKVKLPD-T
AhSRK20        YCDTSTSPMCNCIRGLNPWSMQQ-WDLKDGSSGCVRRTPLSC-SGDGFVPLKKMKLPD-T
AhSRK28        FCDTNTSPNCNCIKGFEPRNPQG-GALENRSTECVRKTQLNC-NGDGFFWLRNMKLPD-T
AhSRK32        YCNTNTAPLCNCIRGFIPWSMQQ-WNLGDGSSGCVRRTPLSC-SGDGFLLMKKMKLPT-T
AhSRK43        YCDMITSPICNCIKGFTPRYSEA-WKLKDGASGCVRKTPVSCNGKDEFVQLKNMKLPD-T
AhSRK47        YCDINTSPPCNCIKGFDPKYPQQ-WELSNGVGGCVRKTRLSC-NDDGFVRLKKMKLPV-T
CgSRK1SD       FCDLNTTPKCNCIKGFEPQG----GTMDNRSTECVRKTPLEC-NGDGFFGLKNMKLPY-T
CgSRK2SD       YCDTNTSPMCNCIRGFRPKFPQA-WILRDGSSGCVRKTRLSC-GRDRFVQLNNMKMPD-T
CgSRK3SD       YCDINTSPNCNCIQGFVPKYPE--WKLIDGAGGCVRRIPLDC-RKDRFLPLKQTKLPD-T
CgSRK4SD       YCDPNERPYCNCIKGFEPRS----GALDNTYTECIRKTQLRC-NGDGFFWLRNMKLPD-T
CgSRK5SD       YCDMNTSPRCNCINGFGPKNPHK-WLLEGGIGECVRKTQLSC-RGDKFVQLKNMKLPD-S
CgSRK6SD       VCDTNTTPSCNCIKGFE---LEG-GTWDSVSSECVRNTQLNC-NGDGFSLLKNMKMPY-T
CgSRK7         YCDMSTSPMCNCIEGFAPRNSQE-WASGIVRGRCQRKTQLSC-GGDRFIQLKKVKLPD-T
CgSRK9SD       YCDSTKTQKCNCIKGFEPMDPRE-GALDNTFTECVRKTQLSC-VDDGFFGLRNMKLPD-T
CgSRK11SD      FCDLSTSPACNCIRGFQPLFPQE-WALGDVRGRCRRKTQLNC-GGDKFVHLMNMKLPD-T
CgSRK18SD      YCDVSTSPECNCIEGFQPPYPQD-WALGDVTGRCQRKKKLNC-TGDKFIRLSNMKLPP-T
Cg2-2-KS2      YCDVNTSPRCNCIQGFDPKNQQQ-WDLSNGVSGCVRRTRLSC-REKRFLRLKKMKLPV-T
CgSRK34SD      YCDTNTKPICNCIEGFEPTNSQE-GALDNTVTECVRKTQSSC-NGDGFFWMKKMKLPS-T
CgSRK35        YCDMDTSPVCNCIQGFTPRSLQD-WVLRDGSNGCVRKTPLSC-GGDGFVLLKKMKLPD-T
CgSRK36SD      YCDVNTSPICNCIPGFNPSDVQQ-WDQRSWSGGCIRRTQLSC-SEDGFTKMKNMKLPE-I
CgSRK37SD      YCYLNTSAFCSCIKGFEPKIQEA-WAVNDGTSGCVRKTRLSCTSGDGFFKLKNTKLPD-T
CgSRK47SD      YCDVNASPVCNCIQGFKPVNLKQ-WDLKTWAGGCMRKTRLSC-SGDGFTRMKNMKLPE-T
```

```
AlSRKb(S-20)  SAAIVDRTIDLGECKKRCLNDCNCTAYASTDIQNGGLGCVIWIE
AlSRK01       KTVIVDRKIGMKDCKKRCLNDCNCTAYANTDI--GGTGCVMWIG
AlSRK04       TMATVDRRISGKECKQKCLMDCNCTAYANADIKNGGLGCVIWTG
AlSRK05       TTAIVDKRIGLKECEEKCKQYCNCTAYANTDTRNGGTGCVIWIG
AlSRK06       TGAIVDKRIGVKECEEKCINNCNCTAFANTNIQDGGSGCVIWTS
AlSRK09       IVATVDRGIGLKECEERCLNDCNCTSFANADVQNGGWGCVIWTG
AlSRK13       TAAIVDKRIGFKDCKERCAKTCNCTAFANTDIRNGGSGCVIWIG
AlSRK14       SGAIVDKRIGLKECEERCIKECNCTAFSNMNIQDGGKGCVIWTK
AlSRK08       MGAIVDKRIGLKECEERCINDCNCTAFANTNIQDRGSGCVIWTG
AlSRK15       EAIVVDRRIGIKECEKRCQIDCNCTAFANIDIQNGGLGCVFWTK
AlSRK16       TTATVDKRLGLEECEQKCKNDCNCTAFANMDIRNGGPGCVIWIG
AlSRK18       AGAIVDKRIGLKECEERCIENCNCTAFANTNIQDGGSGCVLWTR
AlSRK19       TMALVDTRIGLKECEVKCLANCNCTAYANMDIRNGGSGCVIWTR
AlSRK22       MSAIVDMEIDEKDCKKRCLSNCNCTGFANADIRNGGSGCVIWTG
AlSRK23       EEVIVDRRISTKECRERCLGDCNCTAFANTDIQNGGWGCVIWTG
AlSRK25       TAAMVDKRIGLKECEKKCKNDCNCTAYAS--ILNGGRGCVIWIG
AlSRK30       TWTIVNKSIDMEECKERCLRDCNCTAYANTDIRNGGSGCVIWTG
AlSRK31       TEAIVDRIIDVKECEDKCINDCNCTAFAN---------------
AlSRK36       TSVTVDRRIGTKECKKRCLNDCNCTAFANADIRNDGSGCVIWTG
AlSRK37       TAAVVDKRIGLKECEEKCKTHCNCTAYANSDVRNGGSGCIIWIG
AlSRK38       TFTIVDRSIDVKECEERCRNNCNCTAFANADIRHGGSGCVIWTG
AlSRK39       TEVTVDRIVGVEECQNRCITDCNCTAFANV-IRNGGLGCVIWTR
AlSRK50       TMAIVDRSISGKECRTKCLRDCNCTAFANADIQNGGSGCVVWTG
AlSRK52       KDAIVDRIIGFKECEKRCLNNCDCTAFANIDSRNGGLGCLIWTG
AhSRK03       SGAIVDKRIGLKECEERCIENCNCTAFANTNIQDGGSGCVLWTR
AhSRK12       MDAIVDRKIGKKECKERCLGDCNCTAYANID----GSGCLIWTG
AhSRK13       TTAIVDRRLGVKECREKCLNDCNCTAFANADIR--GSGCVIWTG
AhSRK15       KSVIVDKKIDAEECEMRCLQNCNCTAFANKDIRNGGSGCVIWTG
AhSRK20       TMTIVDRRIGVKECKEKCLRNCNCTAYAKADITNGGVGCVIWTG
AhSRK28       SGAIVDKRIGLKECEERCIENCNCTAFANTNIQNGGSGCVLWTR
AhSRK32       TMTTVDRRINWKECREKCLTDCNCTAFANADIQNGGLGCVMWTG
AhSRK43       TSAVVDKRIGLNECRERCLNDCNCTAFANINIQNRGSGCVVWTR
AhSRK47       KDTIVDRRITTKECKKSCLRNCNCTAFANTNIQNGGSGCLIWTG
CgSRK1SD      SGAIVDKSIGLKECEERCSGDCNCTAYANTNLQDGGSGCVMWTS
CgSRK2SD      MQAVLDRRIGAKECRKRCFRDCNCTGFTN--IRNGGWGCVIWTV
CgSRK3SD      KTVIVDRKIGRKDCKKRCLKNCNCTAYANTDI--GGRGCVMWIG
CgSRK4SD      SGAIVDKRIGLKECEDRCIEDCNCTAFANTNVQDGGSGCVLWTS
CgSRK5SD      TGVIVDRRIELKECEGRCKINCNCTAYANTDIQNGGSGCVIWTS
CgSRK6SD      KGAIVDKRIGLKECGKRCIEDCNCTAYANTNIQDGGSGCVMWTS
CgSRK7        TEAIVDKRLGLEDCKKRCATNCNCTAYATMDIRNGGLGCVIWIG
CgSRK9SD      SGAIVDKRIGLKECEDMC--------------------------
CgSRK11SD     TTAIVDKRIGLKECRMKC--------------------------
CgSRK18SD     TEVIVDKTIGIKDCEERC--------------------------
Cg2-2-KS2     MDAIVDRNIGKKECKKRCLTNCNCTAYANVD----RSGCLIWTG
CgSRK34SD     MGATVDKSIGLKECEERCMNDCNCTAFANTDIRNGGSGCVIWT-
CgSRK35       TTAIVDRSIDLKECKEICSRNCNCTGFANADIRNGGTGCVIWTG
CgSRK36SD     RMAIVDRSIGLEECKKRCLSDCNCTALANADIRNGGTGCVFWTG
CgSRK37SD     TWTIVDKSIDVEECKKRC--------------------------
CgSRK47SD     TMAIVDRSIDVKECKKRCLSDCKCTAFANADVRNGGTGCVIWTG
```

Fig S4 Yamamoto et al.

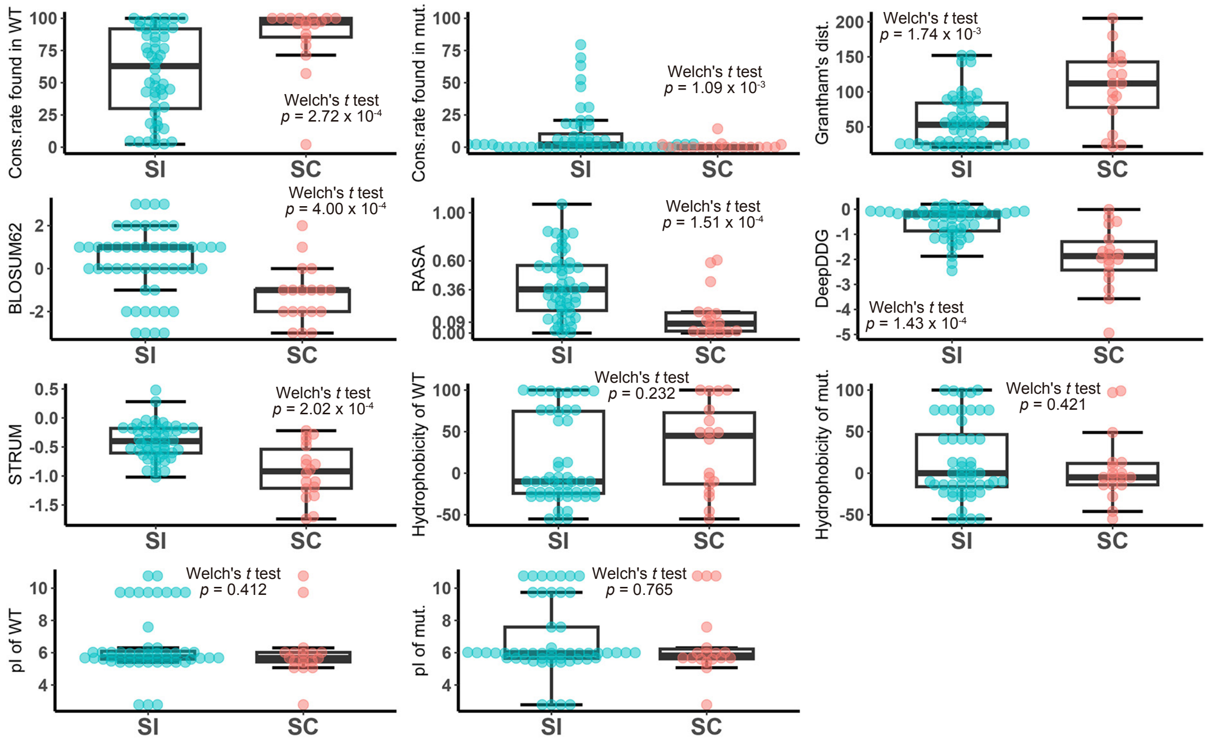


**Supplemental Figure S4.** Comparison of the values of mutations found in *A. thaliana* transformants showing the SC and SI phenotypes.

The values for “Conservation rate (%) of amino acid residues observed in wild-type and mutated AlSRKb proteins across *S* haplotypes” , “Grantham's distance” and “BLOSUM62 score” , “Relative accessible surface area (RASA) of amino acid residues in the AlSRKb structural model” , “DeepDDG score” , “STRUM score” , “Hydrophobicity of amino acid residues observed in wild-type and mutated AlSRKb proteins” , and “pI of amino acid residues in wild-type and mutated AlSRKb proteins” of mutations present in *A. thaliana* transformants showing the SC and SI phenotypes were compared. Welch’ s t tests were performed to determine statistically significant differences between groups.
